# Continued dysfunction of capillary pericytes promotes no-reflow after experimental stroke *in vivo*

**DOI:** 10.1101/2023.03.06.531258

**Authors:** J Shrouder, S Filser, DP Varga, S Besson-Girard, U Mamrak, B Bulut, FB Seker, B Geserich, F Laredo, A Wehn, I Khalin, P Bayer, A Liesz, O Gökce, N Plesnila

## Abstract

Incomplete reperfusion of the microvasculature (“no-reflow”) after ischemic stroke damages salvageable brain tissue. Previous ex-vivo studies suggest pericytes are vulnerable to ischemia and may exacerbate no-reflow, but the viability of pericytes and their association with no-reflow remains underexplored in vivo. Using longitudinal *in vivo* 2-photon single-cell imaging over seven days we show 87% of pericytes constrict during cerebral ischemia, remain constricted post-reperfusion and 50% of the pericyte population are acutely damaged. Moreover, we reveal ischemic pericytes are fundamentally implicated in capillary no-reflow by limiting and arresting blood flow within the first 24 hours post-stroke. Despite sustaining acute membrane damage, we observe up to 80% of cortical pericytes survive ischemia, upregulate unique transcriptomic profiles and replicate. Finally, we demonstrate delayed recovery of capillary diameter by ischemic pericytes after reperfusion predicts vessel reconstriction in the sub-acute phase of stroke. Cumulatively, these findings demonstrate surviving cortical pericytes remain both viable and promising therapeutic targets to counteract no-reflow after ischemic stroke.

## Introduction

Ischemic stroke is an acute-onset disease caused by the occlusion of a large cerebral artery and represents a major cause of death and disability worldwide. Current therapy is restricted to recanalization of the occluded vessel, but incomplete reperfusion of cerebral microvasculature (‘no-reflow’) causes further tissue damage and severely limits recovery in up to 83% of patients post-recanalization ^1–3^.

While the pathogenesis of no-reflow is multi-faceted, a critical step may involve constriction of cerebral microvessels by mural cells such as smooth muscle cells (SMCs) and capillary pericytes, which modulate hemodynamic resistance by controlling capillary diameter in response to neuronal activity ^4–12^. Hybrid SMCs at the arteriole-to-capillary transition are reported to constrict and reduce blood flow to downstream capillaries after ischemia^13^, but whether pericytes themselves have the potential to restrict blood flow within the capillary bed, the vascular territory where no-reflow occurs, remains contentious^14^.

Recent mouse models of global ischemia and experiments in brain slices suggest that pericytes are sensitive to short-term ischemia, contracting around capillaries, sustaining lethal damage, and potentially limiting blood flow long-term ^15, 16^. However, after extended periods of ischemia, longitudinal in vivo evidence for this hypothesis remains sparse, and the few studies that exist disagree about ischemic pericyte contractility ^13–19^. As a result, it is currently unclear whether pericytes survive severe ischemia, how they are implicated in no-reflow development, and whether they represent a viable cellular treatment target within a timescale relevant to stroke patients.

Therefore, in the current study we used repetitive *in vivo* 2-photon microscopy in combination with laser speckle perfusion contrast imaging to dynamically investigate whether pericytes survive ischemic stroke, contract during ischemia and after reperfusion, and are causally implicated in the no-reflow phenomenon.

## Results

### Baseline characterization of capillary pericytes after the arteriole-to-capillary transition

We first characterized cortical capillary pericytes in PDGFRßEGFP pericyte reporter mice with *in vivo* 2-photon imaging and cerebral blood flow using laser speckle contrast imaging (LSCI) in the healthy somatosensory cortex perfused by the distal middle cerebral artery (MCA) (Supplementary Figure 1, Figure 1a). Focusing on previously described distinct capillary pericyte subtypes after the arteriole-to-capillary transition^20^, we found mesh pericytes were present on lower order capillaries (mean 2^nd^ branch order ± 0.8) with an average lumen diameter of 5.2 µm ± 0.9. Junctional thin-strand pericytes at capillary bifurcations were present on a wide range of branch orders (mean 3^rd^ branch order ± 1.4) on capillaries with a lumen diameter of 3.9 µm ± 0.65. ‘En-passant’ thin-strand pericytes which run along capillaries were located deeper within the capillary bed, on higher branch order capillaries (mean 4^th^ branch order ± 1.3) with an average capillary lumen diameter of 4.0 µm ± 0.5 (**Supplementary Figure 2a – c**).

### Pericytes constrict during ischemia and recover slowly during reperfusion

Next, using a protocol with blinded randomization of mice to experimental groups, we confirmed induction of cerebral ischemia via filament middle cerebral artery occlusion (MCAo) produced a 78% reduction in cerebral blood flow (CBF) (**Figure 1b**). CBF remained significantly reduced 90 min after middle cerebral artery (MCA) recanalization and was persistently impaired seven days after ischemia, indicating incomplete microvascular reperfusion (**Figure 1c**). Within the first week of stroke, MCAo resulted in the loss of neurons, the development of an ischemic lesion, and significant neurological deficits (**Supplementary Figure 3a – f**) ^20^.

**Fig. 1.**
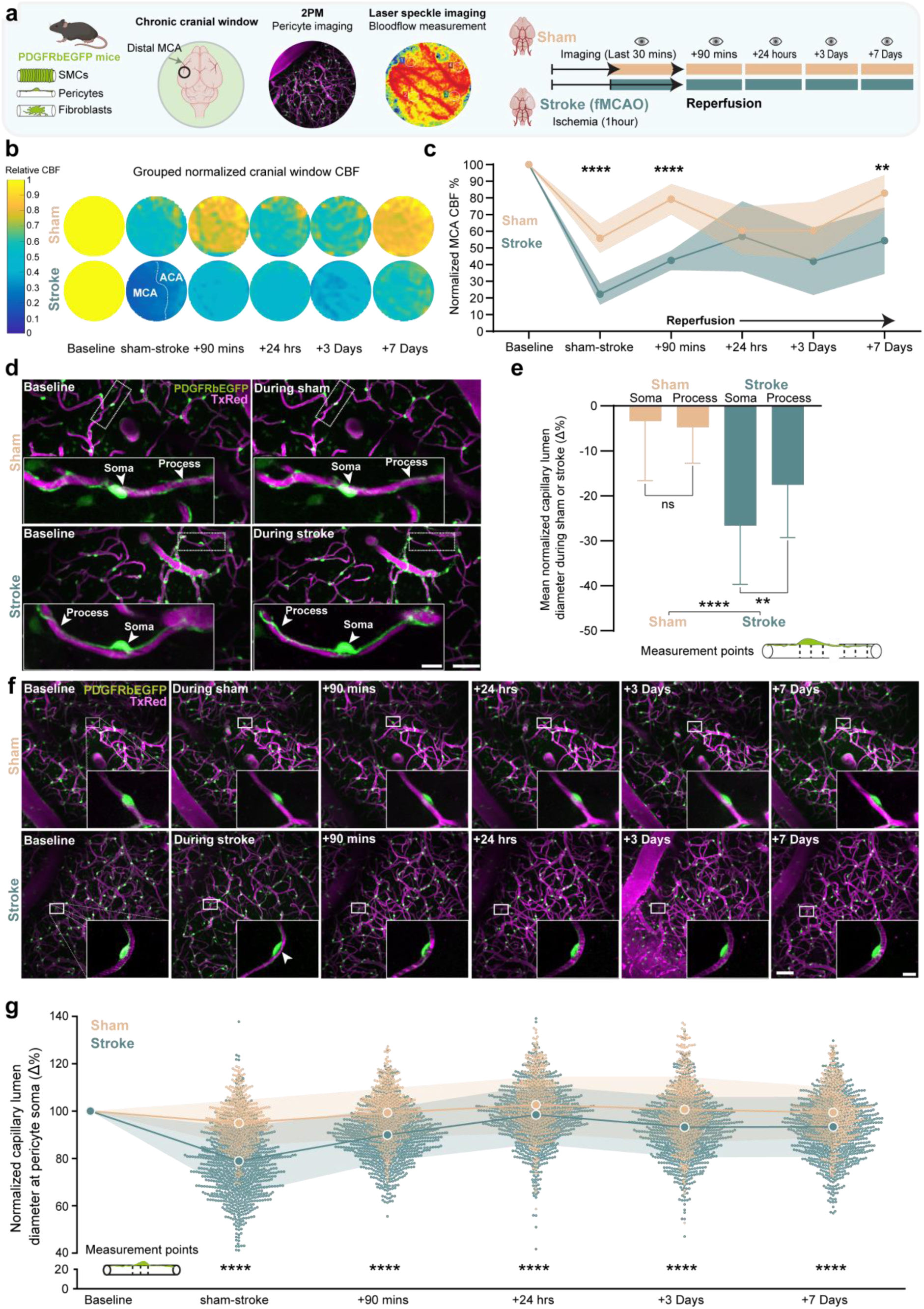
Pericytes constrict in response to transient experimental ischemia in vivo. **a)** In vivo multi-modal imaging methodology, experimental groups and imaging timeline in PDGFRßEGFP mice. **b)** Grouped normalized LSCI CBF perfusion within the chronic cranial window. **c)** Normalized LSCI CBF within the territory perfused by the distal MCA per experimental group. **d)** 2-PM z-projections of capillary beds imaged at baseline and during stroke or sham surgery, blood vessels are shown in magenta, while mural cells are shown in green. Inserts, individual capillary pericytes with white arrows highlighting pericyte soma and processes. **e)** Quantification of mean normalized capillary lumen diameter changes at pericyte soma or processes during either stroke and sham surgery relative to baseline. **f)** Upper, 2-PM z-projections of fields of view imaged over experimental series. Inserts, zoomed in 2-PM imaging of individual pericytes over experimental timeline. **g)** Mean normalized lumen diameter change at pericyte soma over experimental time course between stroke and sham groups respectively, individual pericytes shown in blue (stroke) and gold (sham). n=7 stroke mice and 4 sham mice ∼1400 pericytes stroke-890 sham-510 repeatedly measured across 6 time points. Statistics **c,** Two-way ANOVA with Greenhouse-Geisser correction. e, Unpaired Student’s t-test was used to compare sham against stroke pericyte measurements and Paired two-tailed Student’s t-test was used to compare 120 pericyte soma vs process measurements against baseline in 5 stroke mice and 124 pericyte soma vs process measurements in 4 sham mice. **g** One-way ANOVA, Bonferroni Post-hoc correction and pairwise two-tailed Student’s t-test. In each figure, significance codes *p < 0.05, **p < < 0.01, ***p < 0.001 and ****p < 0.0001. Scale bars **d,** 50 µm and 10 µm, **f,** 50 µm and 10 µm. Data is shown as mean +/-s.d and source data are provided as a source data file.

To determine if ischemia led to a decrease in vessel lumen diameter due to pericyte constriction *in vivo*, we measured capillary lumen diameter at both the cell soma and processes in randomly selected ‘En-passant’ thin-strand pericytes during either sham or stroke surgery (**Figure 1d**). We found ischemia reduced capillary lumen diameter at both pericyte soma and processes compared to baseline or sham surgery (**Figure 1d**). Specifically during ischemia, capillary lumen diameter was significantly decreased at pericyte soma compared to pericyte processes (27% ± 13% vs 18% ± 11%), while no differences were found between soma and process measurement during sham surgery or when compared to baseline (**Figure 1e**). Thus, even in capillary pericytes suggested to express a limited pool of contractile machinery^7^, these data suggest ischemic pericytes exert substantial contractile force focally at the cell soma and are also capable of imparting contractile tone at their processes, consistent with recent optogenetic probing of pericyte contractility ^10, 19^.

To understand whether pericytes may mediate incomplete reperfusion after MCAo at a population level, we next followed about 900 individual capillary pericytes during and after cerebral ischemia over a period of seven days to assess their influence on vessel diameter compared with more than 500 sham pericytes (**Figure 1f**). During stroke, 87% of all investigated capillary pericytes constricted, with the average capillary lumen diameter decreasing by 21 ± 14% of baseline beneath the pericyte soma and the maximum pericyte constriction reaching 60% of baseline (**Figure 1g**). Importantly, 90 minutes after reperfusion of the occluded vessel, pericytes still significantly constricted their associated capillary by 11 ± 11%. At 24 hours post stroke, pericytes dilated to a degree approaching pre-stroke baseline levels, but remained significantly constricted compared to pericytes in sham-operated mice. Interestingly, pericytes in the reperfused brain entered a secondary phase of constriction on days three and seven post-stroke, constricting by 7 ± 13% compared to sham pericytes. Cumulatively, these data suggest that according to the Hagen-Poiseuille law pericytes severely reduce cerebral blood flow during and acutely after ischemia by up to 87% and may thus be involved in no-reflow development. Moreover, the heterogeneity in constriction severity suggested not all pericytes constrict to the same degree; therefore, we next sought to dissect the contractile properties of ischemic pericytes subtypes.

### Pericyte subtypes constrict uniquely during and after ischemia

Capillary pericytes have recently been characterized into subtypes based on their morphology and location within the microvasculature ^20^. During ischemia, we observed a longitudinal biconcave morphology of constriction in the mesh pericyte population, while junctional and en-passant thin-strand pericytes appeared to constrict focally near the cell soma (**Figure 2a, white arrows**). This constriction was evident at a population level, where all subtypes constricted heavily during stroke and acutely post reperfusion compared to their sham counterparts (**Figure 2b, c**). Specifically, en passant thin-strand pericytes constricted the most during ischemia (**Figure 2c**, constriction of en-passant thin-strand pericytes: 25% vs junctional:21% p<0.0001, or mesh: 20% p<0.0005). Moreover, we observed a cortical depth dependent constriction severity in thin-strand pericytes that was absent from mesh pericytes (**Figure 2d, e left**). Specifically, en passant thin-strand pericytes between 50-100 µm below the brain surface constricted significantly less than those 150-200 µm below the surface (**Figure 2e, middle**), while junctional thin-strand pericytes 50-100 µm below the surface constricted significantly less than those 200-250 µm below the surface (**Figure 2e, right**).

**Fig. 2.**
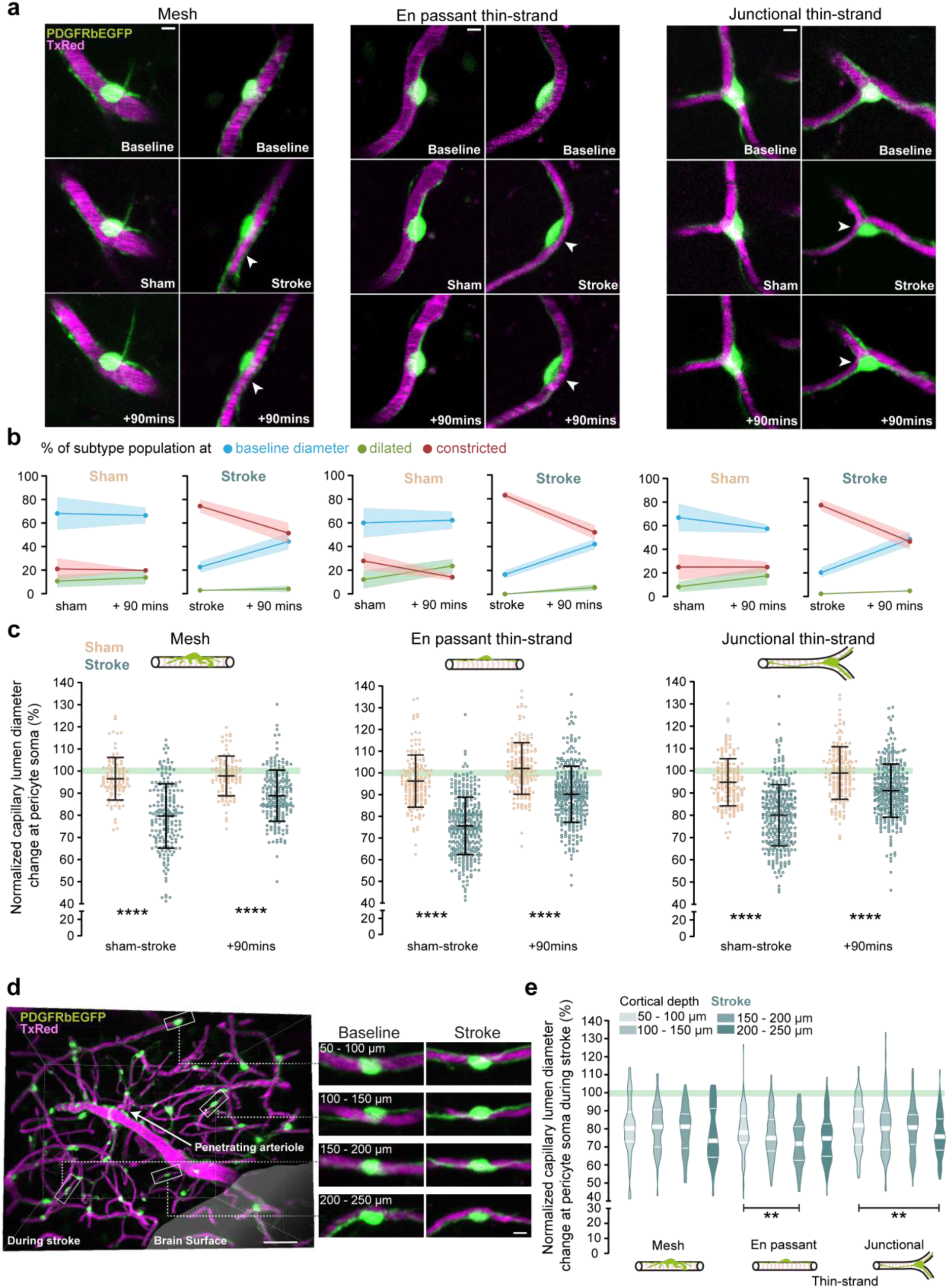
Ischemic pericytes constrict in a subtype and depth dependent fashion during stroke and remain constricted acutely post reperfusion. **a)** 2-PM imaging of each pericyte subtype at baseline, during stroke/sham and 90 mins post-surgery. **b)** Percentage of each subtype population per group either constricting by more than 10% (re**d)** dilating by more than 10% (gree**n)** or within 10% of baseline diameter (purple). **c)** XY graphs of normalized individual pericyte subtypes during stroke/sham surgery, 90 mins post reperfusion. **d)** Imaris® 3D view of a single penetrating arteriole and constriction of en passant thin-strand pericytes at differing cortical depths. **e)** Quantification of individual pericyte constriction during stroke per cortical depth, per subtype analyzed. n=7 stroke, 4 sham PDGFRßEGFP mice. Statistics, **a,** lower, two-way ANOVA with Greenhouse-Geisser correction. **c, e** unpaired two-tailed Student’s t-test. In each figure, significance codes *p < 0.05, **p < < 0.01, ***p < 0.001 and ****p < 0.0001. Scale bars **a,** 5µm, **c,** 50 µm and 5 µm. Data is shown as mean +/-s.d and source data are provided as a source data file.

After establishing that pericytes remain contracted after ischemia in a subtype and cortical depth dependent fashion, we next asked if pericyte constriction led to interruptions in microvascular flow.

### Blood flow is arrested at pericyte soma during and after ischemia

Before cerebral ischemia microvascular structure and flow was complete and contiguous, with plasma dye labelling the complete capillary bed from penetrating arteriole to ascending venule (**Figure 3a, Baseline**). However, during stroke multiple regions of the cerebral microvasculature were free of plasma, indicating a complete cessation of flow (**Figure 3a, white arrows**). We subsequently quantified the prevalence of these capillary stalls as a function of cortical depth. We observed significantly more capillary stalls in the microvasculature during stroke and 90 minutes post reperfusion than after sham surgery across all cortical depths measured. Notably, most capillary stalls were located in superficial cortical layers after the arteriole-to-capillary transition, suggesting capillary pericytes may be involved in arresting blood flow (**Figure 3b**). Therefore, we next asked whether pericyte soma were proximally associated with capillary stalls (**Supplementary Figure 4a**). We found pericytes were associated with capillary stalls during both sham and stroke surgery (**Figure 3c, white arrows**), however almost eight times more pericytes were associated with capillary stalls during stroke than after sham surgery (31 ± 8% vs 4 ± 2%). Importantly, this association continued 90 minutes post reperfusion, where 6 ± 5% of the ischemic pericyte population remained associated with capillary stalls (**Figure 3d**). Furthermore, significantly more ischemic pericytes were associated with stalls up to 24 hours post-stroke, where they consistently accounted for one third of all capillary stalls in the ischemic brain (**Figure 3e, Supplementary Figure 4b**). These data indicate that pericytes are readily associated with capillary stalls under conditions of ischemia and demonstrate that pericytes participate in the ‘no-reflow’ phenomenon *in vivo* via cessation of blood flow proximal to the cell soma.

**Fig. 3.**
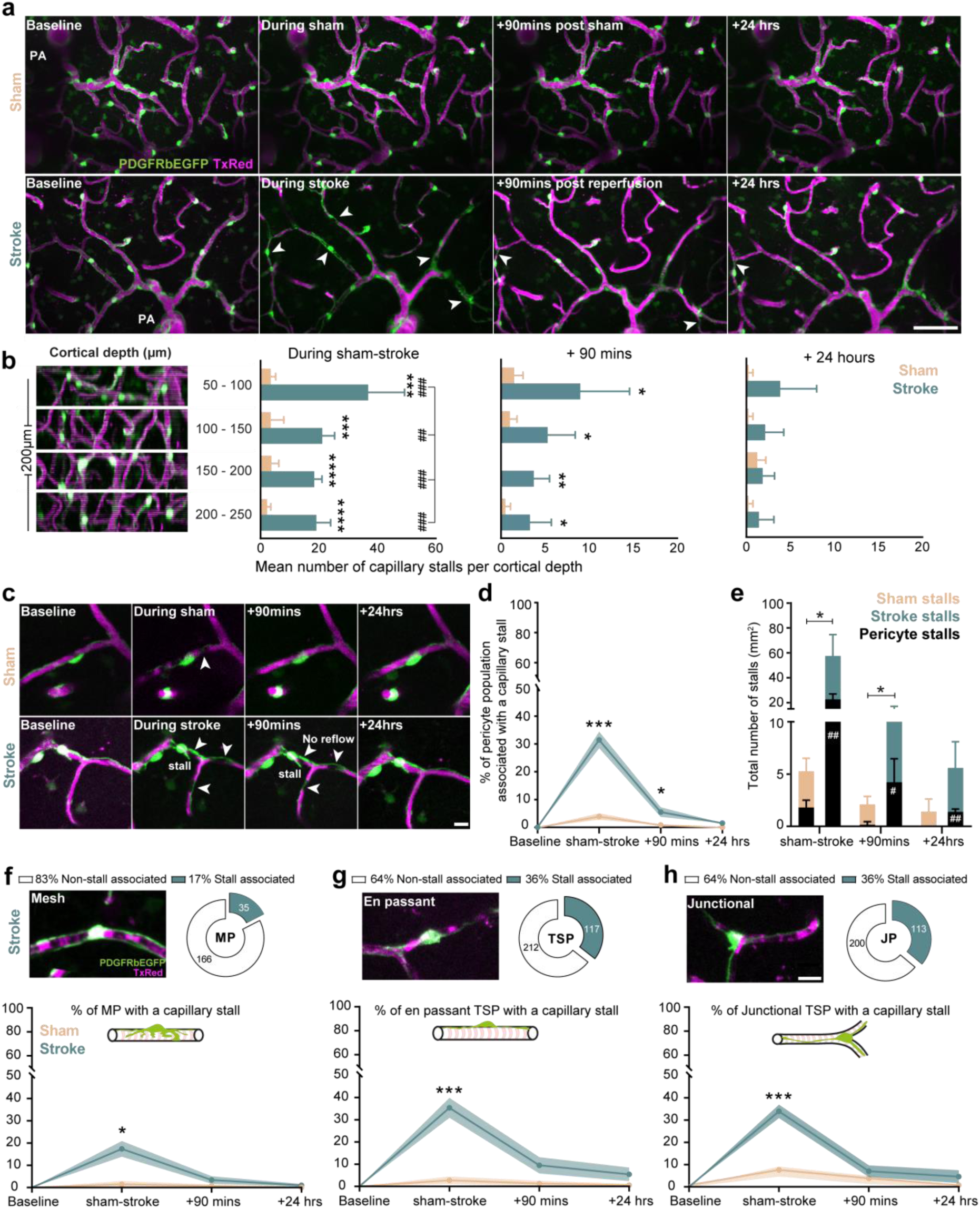
Ischemia induces capillary stalls at pericyte locations during stroke and acutely after reperfusion. **a)** Left, capillary stalls during either stroke or sham surgery, white arrows indicate sites of capillary stalls **b)** Quantification of mean capillary stalls per cortical depth/ROI during either stroke or sham surgery. **c)** 2-photon imaging of capillary stalls during and after sham or stroke surgery. **d)** Percentage of capillary stalls associated with either stroke or sham pericytes over time. **e)** Mean total number of capillary stalls per mm^2^ in either sham or stroke group either pericyte associated or non pericyte associated. **f, g, h)** Upper, percentage of each pericyte subtype associated with capillary stalls during stroke. Lower, percentage of capillary stalls associated with either stroke or sham pericyte subtypes over time. n=7 stroke and 4 Sham mice. Statistics, **b**, unpaired Welch’s t-test, within groups one-way ANOVA **d**, **e**, **f**, **g**-Two-way ANOVA with Sidak’s multiple comparisons test. In each figure, significance codes *p < 0.05, **p < < 0.01, ***p < 0.001 and ****p < 0.0001. Scale bars, **a,** 50 µm **c,** 10 µm and **f,g,h,** 10 µm respectively. Data is shown as mean +/-s.d for **b**, for **d**, **e**, **f** – mean +/-s.e.m and source data are provided as a source data file.

We next evaluated which pericyte subtype populations were most frequently associated with capillary stalls during stroke. All pericyte subtypes showed a significantly higher association with a capillary stall after stroke compared to sham surgery, but this association was significantly more prevalent in en passant and junctional thin-strand pericytes compared to mesh pericytes (**Figure 3f-h, upper panels and lower graphs**, **Supplementary Figure 4c**). Collectively, these data suggest that thin-strand pericytes predominantly impair microvascular flow during ischemic stroke and in the first 24 hours after reperfusion. In the next step, we investigated the possible reasons for pericyte-associated flow disturbances after cerebral ischemia.

### Ischemia damages pericytes and cellular stress remains visible acutely post-reperfusion

Together with observations of pericyte contractility, *in vivo* 2-photon imaging revealed blebbing in pericytes subjected to cerebral ischemia (**Supplementary Figure 5a, b**). These findings mirrored Rho-kinase dependent pericyte blebbing reported during retinal ischemia^18^, and we therefore asked whether cytoplasmic alterations were visible in pericytes 90 minutes post reperfusion in PDGFRßEGFP mice *ex vivo*. The ischemic region demarcated by the loss of Iba1^+^ microglia/macrophages and GFAP^+^ astrocytes contained EGFP^+^ pericytes of varying fluorescence intensity and segregated clearly from EGFP^+^ pericytes in ipsilateral tissue, where the signal remained bright and uniform (**Figure 4a, inserts, 1, 2, 3**). We then used Aquaporin-4 to label astrocytic end-feet and collagen IV staining to label the basement membrane in which the entire pericyte cytoplasm is contained under physiological conditions. Here, with super-resolution confocal microscopy, we observed EGFP^+^ pericytes split into two phenotypes: pericytes with endogenous EGFP labelling the entire cytoplasm within the basement membrane and pericytes with cytoplasmic shedding of EGFP beyond the basement membrane (**Figure 4b, c white arrows**). After verifying these morphological signs of pericyte damage using line profile measurement (**Supplementary Figure 5c**), we quantified the distinct phenotypes and observed that 57% of the mesh pericyte population, 59% of en passant thin-strand pericytes and 48% of junctional thin-strand pericytes appeared to have suffered membrane damage within the infarct area (**Figure 4d, e, f, middle right and Imaris^®^ 3D reconstructions**). To test for potential cytoplasmic leakage as a result of membrane damage. We then used nuclear staining (DAPI) to compare EGFP fluorescence intensity within the nuclei of each pericyte phenotype (**Figure 4d, e, f, middle**). We observed no difference between damaged and intact mesh pericytes within the infarct. However, damaged infarct core mesh pericyte nuclei contained significantly less EGFP intensity compared to contralateral mesh pericytes (**Figure 4d**). In contrast, damaged en passant thin-strand and junctional thin-strand pericytes had significantly less EGFP intensity in their nuclei than those with an intact cytoplasm within the infarct, while intact pericytes had the same EGFP intensity as those in contralateral hemisphere (**Figure 4e, f**). Cumulatively, these data indicate pericytes incur substantial damage-associated morphological alterations during ischemia and acutely after reperfusion, blebbing and shedding portions of their cytoplasm in the absence of blood flow. Furthermore, decreases in nuclear EGFP intensity within damaged, but not intact pericytes suggested capillary pericyte membrane integrity may be critically compromised after ischemia, therefore, we next sought to determine whether these signs of cellular stress resulted in broad scale pericyte death.

**Fig. 4.**
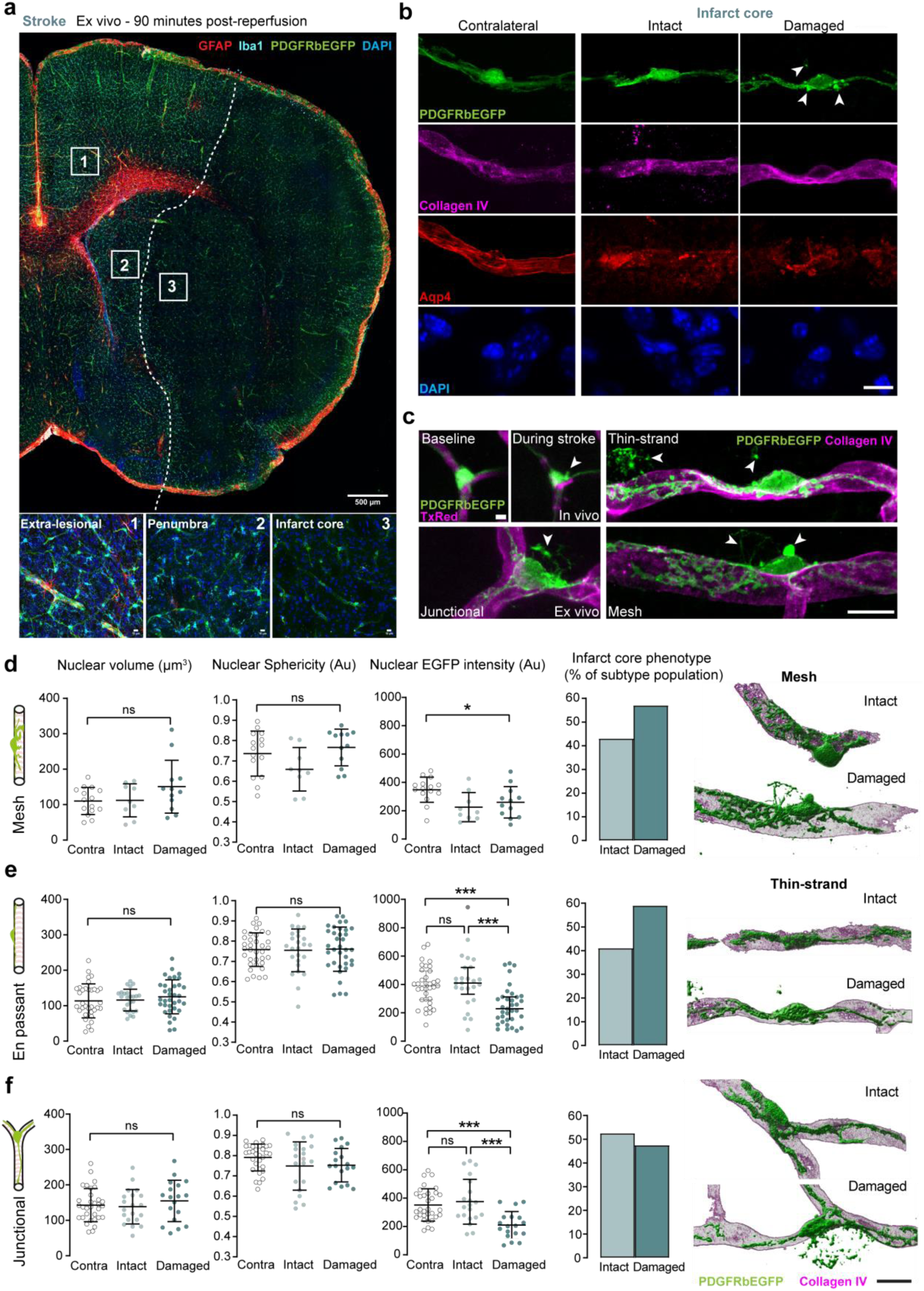
Ischemia damages pericytes and cellular stress remains visible acutely post-reperfusion. **a)** Upper, *Ex vivo* brain slice overview in a PDGFRßEGFP mouse 90 mins post reperfusion, staining for microglia (iba1) shown in cyan, astrocytes (GFAP) in red, nuclear (DAPI) staining in blue. **b)** Super-resolution airyscan imaging of PDGFRßEGFP^+^ pericyte phenotypes within the infarct core, basement membrane component collagen IV (magenta), Aqp4^+^ labelling astrocytic end-feet (red), with DAPI^+^ nuclear staining (blue). **c)** Upper left, 2-PM In vivo imaging of pericyte blebbing during ischemia. Lower left and Right, Ex vivo airyscan imaging of damaged pericyte subtypes within ischemic territory. **d, e, f)** Left, nuclear sphericity, volume and nuclear EGFP intensity of intact, damaged and contralateral pericyte subtypes. Middle right, percentage of intact vs damaged pericytes. Right, Imaris 3D reconstruction and proportion of intact/damaged pericyte phenotypes within the infarct core.. n=4 stroked PDGFRßEGFP mice, Statistics, **f** Kruskall-Wallis test (Nuclear EGFP intensity) and **f lower,** unpaired two-tailed Student’s t-test. In each figure, significance codes *p < 0.05, **p < < 0.01, ***p < 0.001 and ****p < 0.0001. Scale bars, **a,** 500 µm and 10 µm, **b, c,** 5 µm and 10 µm **d, e, f,** 10 µm. Data is shown as mean +/-s.d, for F (Nuclear EGFP intensity)-Median with interquartile range and source data are provided as a source data file.

### The majority of pericytes survive transient cerebral ischemia

During longitudinal *in vivo* imaging, we found a gradual loss in visibility of ischemic pericytes in deeper cortical layers compared to pericytes in sham operated mice (**Figure 5a, Supplementary Figure 6, 11**). Specifically, in mice subjected to cerebral ischemia, the number of pericytes visible by 2-photon microscopy decreased significantly 24 hours after reperfusion and reached a 21 ± 27% reduction by day 7 post-stroke (**Figure 5b, upper, lower**). To assess whether the loss of pericyte visibility may have occurred because of pericyte cell death, we subjected adult male C57Bl/6J mice to sham or stroke surgery (**Supplementary Figure 7**) and assessed pericyte death using TUNEL staining and PDGFRb antibodies to label pericytes (**Figure 5c**). One and three days post-stroke TUNEL^+^ pericytes were identified within infarct core regions, while we found no evidence of TUNEL^+^ pericytes within the sham group (**Figure 5c**). Pericyte density in the infarct core and peri-infarct regions was reduced by ∼ 25% compared to contralateral regions after 24 hours, and by 16% and 15% respectively, on day three post stroke (**Figure 5d**). We then quantified the number of TUNEL^+^ PDGFRß^+^ cells (**Figure 5d, red bars**) and found that stroke induced the appearance of significantly more TUNEL^+^ pericytes in both infarct core and peri-infarct regions compared to the contralateral hemisphere 24 hours post-stroke (**Figure 5e**). However, on day three post stroke only 3% of pericytes within the infarct core and peri-infarct regions were TUNEL^+^ while no TUNEL^+^ pericytes were observed in the contralateral hemisphere (**Figure 5e**). Moreover, we found significantly more TUNEL^+^ pericytes were present on day one post-stroke than day three post-stroke within infarct and peri-infarct regions. Taken together, our data suggest that transient cerebral ischemia results in a modest amount of acute pericyte cell death and demonstrates that pericytes are surprisingly resilient to transient ischemia.

**Fig. 5.**
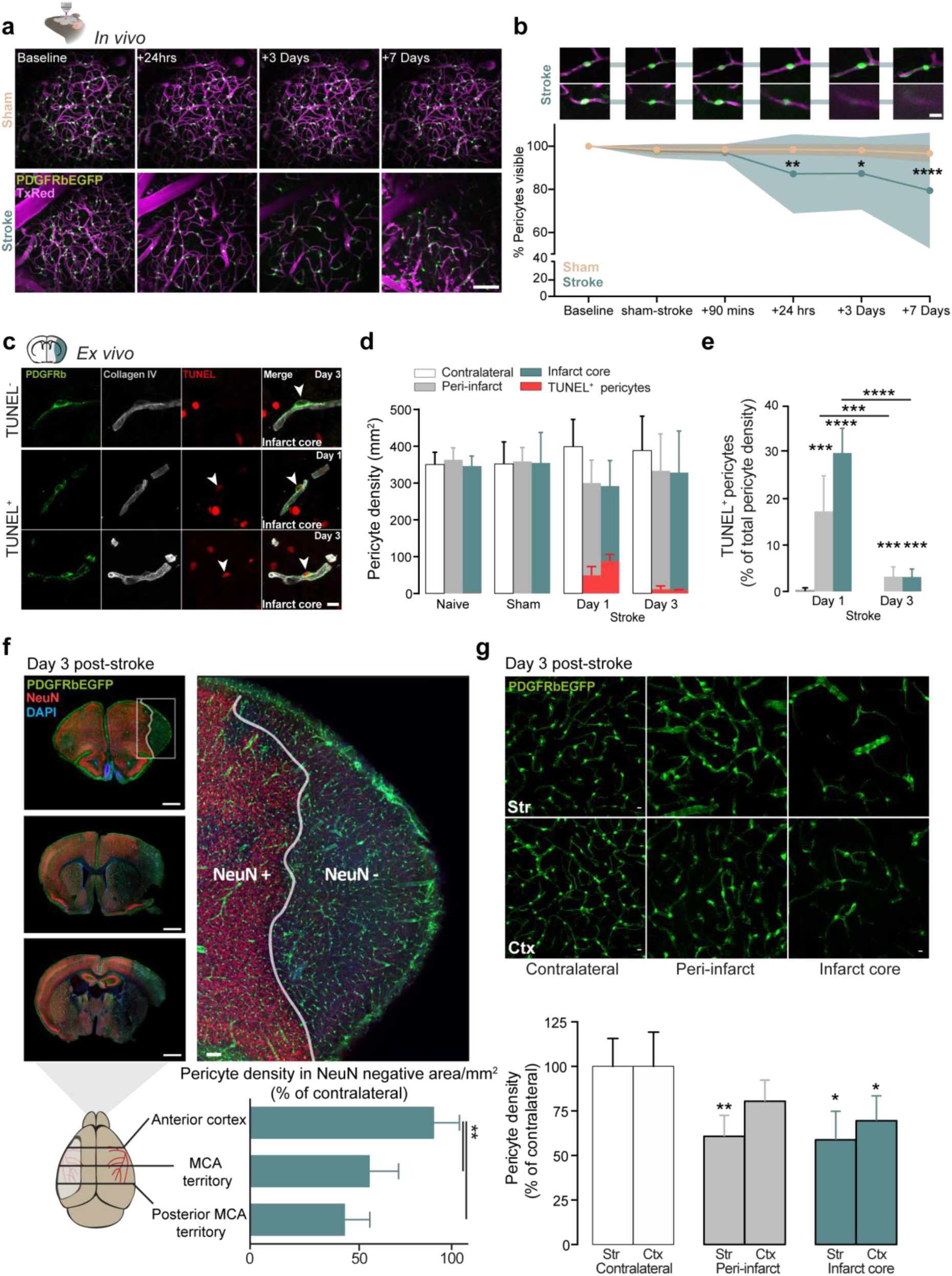
Pericyte cell death peaks acutely after ischemia but the majority of pericytes survive in the infarct. **a)** 2-PM imaging of capillary beds in either sham or stroke group, individual EGFP^+^ pericytes are shown in green, vascular lumen is labelled with TexasRed (Magenta). Note the gradual loss in pericyte visibility over time in the stroke group. **b)** Upper, individual examples of capillary pericytes within the stroke group which remain visible after ischemia, or are lost from their associated vessel. Lower, percentage of pericytes remaining visible after either sham or stroke surgery. **d)** Pericyte density per mm^2^ in each region, number of TUNEL^+^ pericytes are highlighted in red. **e)** TUNEL^+^ pericytes expressed as a percentage of the total pericyte density recorded on day 1 or day 3 post-stroke. **f)** Upper, caudal-rostral brain slice overviews of PDGFRßEGFP, NeuN and DAPI on day 3 post-stroke. Lower, caudal-rostral pericyte density within the region of neuronal loss expressed as a percentage of the contralateral pericyte density. **g)** Upper, confocal overviews of EGFP^+^ pericytes in striatum or cortex 3 days post-stroke. Lower, pericyte density in each respective region as a percentage of the same region in the contralateral hemisphere. Statistics **b** two-way ANOVA with Greenhouse-Geisser correction, **e,f,g** Unpaired Student’s t-test. In each figure, significance codes *p < 0.05, **p < < 0.01, ***p < 0.001 and ****p < 0.0001. **a, b** n=7 stroke and 4 Sham PDGFRßEGFP mice, **c, d, e** n=5 male C57BL/6J mice per group **f, g** n=4 PDGFRßEGFP mice per quantification. Scale bars, **a,** 50 µm, **b, c, g,** 10 µm, **f,** 1 mm and 50 µm. Data is shown as mean +/-s.d and source data are provided as a source data file.

### Pericyte survival is region dependent

To understand if pericytes are more susceptible to cerebral ischemia than other cell populations and whether their survival depends on anatomical location, adult PDGFRßEGFP mice were sacrificed on day three after cerebral ischemia and stained for neuronal, astrocytic, and microglial markers (NeuN, GFAP/Aqp4, Iba1). A qualitative assessment of astrocytes (**Supplementary Figure 8**) and neurons indicated that both cell types were eliminated from the infarct core by day three, while the majority of pericytes remained visible (**Figure 5f**). EGFP^+^ pericytes could even be visualised within the infarct core where all viable neurons had died and revealed a caudal-rostral axis dependency of survival (**Figure 5f, left panels**). Pericyte loss was highest within the posterior stroked MCA territory, with a highly significant reduction in pericyte cell density of 56%. Proximal to the infarcted MCA territory, pericyte density was significantly reduced by 45%. In the anterior infarct core cortical pericyte density was reduced by only 15%, leaving 85% of pericytes viable (p>0.05, compared to contralateral anterior cortex, **Figure 5f, lower panel**). These results suggest pericyte survival may depend on collateral perfusion and, hence, the degree of cerebral ischemia, but that pericytes are less susceptible to cerebral ischemia than neurons. To assess pericyte survival in infarct and peri-infarct areas of the brain, we examined coronal sections taken from the MCA territory of PDGFRßEGFP mice on day three post stroke (**Figure 5g**). Pericyte density was significantly reduced in both infarct and the peri-infarct regions, but differences in pericyte density between cortical and striatal regions did not reach significance (**Figure 5g**). Cumulatively, these data reveal that the majority of ischemic pericytes survive transient cerebral ischemia long-term in a region dependent manner.

### Surviving pericytes enter the cell cycle and replicate

Our data on pericyte density after cerebral ischemia indicate that despite broad-scale neuronal cell death and modest pericyte loss, vessel density remains constant and pericyte coverage increases after stroke (**Supplementary Figure 9a, b, c**). These findings suggested surviving pericytes may be responding to stroke in a compensatory manner. Therefore, we isolated EGFP^+^ pericytes from the infarct core, ipsilateral and contralateral brain regions three days post-stroke using FACS and performed a region specific transcriptomic analysis (**Figure 6a, Supplementary Figure 10a**)^21^. To verify isolated EGFP^+^ cells are indeed pericytes, we compared the transcriptomic profile against a previously defined pericyte signature and found pericyte transcripts to be highly enriched within our dataset (*PDGFRß, Cspg4, Kcnj8, Rgs5, Abcc9*, **Supplementary Figure 10c,d, Supplementary Table 1**)^22^.

**Fig. 6.**
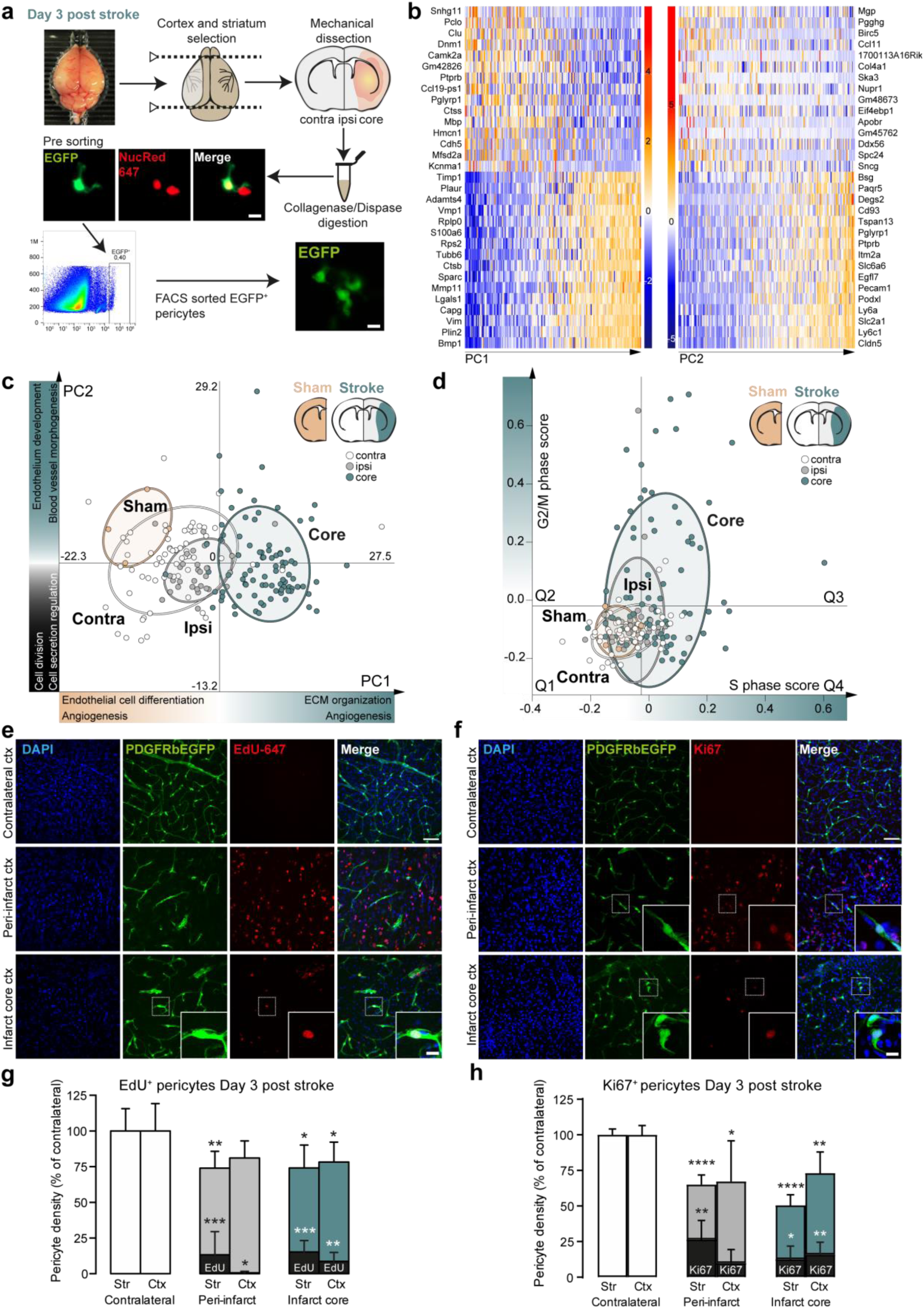
Surviving pericytes enter the cell cycle and replicate where cell density is reduced. **a)** FACS isolation workflow of pericyte isolation day 3 post stroke **b)** Differentially expressed genes contributing the most to variation in core samples relative to contralateral samples on Day 3 post stroke. Data is shown as z-score. **c)** Principle component analysis of region specific sample variation in Day 3 post-stroke brain samples: core, ipsi, contra relative to sham animals. **d)** Average gene expression scores of region specific sample variation in cell cycle associated transcripts on Day 3 post-stroke brain samples relative to sham animals. Quadrants were delimited by the maximum values for G2/M and S-phase score. **e)** Region specific images of EGFP^+^ cells stained with DAPI (blue) and EdU647 (red). **f)** Region specific images of EGFP^+^ cells stained with DAPI (blue) and Ki67 (red). **g)** Quantification of pericyte density and percentage of EdU^+^ pericytes on day 3 post stroke.**h)** Quantification of pericyte density and percentage of Ki67^+^ pericytes on day 3 post stroke.. n=4 PDGFRßEGFP mice per quantification, Statistics, **g, h** unpaired two-tailed Student’s t-test. In each figure, significance codes *p < 0.05, **p < < 0.01, ***p < 0.001 and ****p < 0.0001. Scale bars **a,** 10 µm, **b,** 1 mm, **c, f,** 25 µm, **d, g,** 10 µm. Data is shown as mean +/-s.d and source data are provided as a source data file.

Using principal component analysis, we distinguished few genes were contributing to the transcriptomic variance in distinct brain regions and performed gene ontology analyses on the altered transcriptomic states (**Figure 6b**). We found that PC1^-ive^ samples were associated with endothelial cell differentiation (**Figure 6c**,-log10p6.247), while PC1^+ve^ samples from the infarct core were associated with extracellular matrix reorganisation (**Figure 6c**,-log10p16.74). PC2^+ve^ samples from contralateral and sham brain regions associated with blood vessel morphogenesis pathways (-log10p-15). Interestingly, PC2^-ive^ samples of the stroked brain, regardless of region were dominantly associated with cell division (-log10p-6.5). During previous histological analyses, we observed the appearance of Collagen IV^+^/PDGFRß^+^ cell clusters suggestive of proliferation post stroke (**Supplementary Figure 9d**). Therefore, we tested whether pericytes expressed transcripts associated with cell cycle entry (**Figure 6d**)^23^. We found sham samples expressed low/negative average scores for cell cycle entry (**Supplementary Table 2, Quadrant 1, Figure 6d**), indicating pericytes from ipsilateral sham tissue likely remain in interphase. However, 3 days post-ischemia in the stroked brain, bulk isolated pericytes from all regions radiated into positive gene expression quadrants associated with G2/M phase gene expression (Quadrant 2), G2/M phase and S-phase associated gene expression (Quadrant 3) or S-phase associated gene expression (Quadrant 4). Specifically, 22% of contralateral samples expressed high gene expression scores for S-phase of the cell cycle, and 30% of ipsilateral samples associated with S-phase (Quadrant 4). Within core tissue, we found 11% of samples isolated from the ischemic core were associated with G2/M phase of the cell cycle (Quadrant 2), 28% of samples were associated with both G2/M phase and S-phase phase (Quadrant 3) and 38% of samples were associated primarily with S-phase (Quadrant 4). These data suggest a gradient of cell cycle activation and entry in pericytes recovering from stroke.

To test this at a functional and spatial level, we injected PDGFRßEGFP mice with EdU 24 hours after stroke until sacrifice on day 3 (**Figure 6e**). We observed that 15% of all cells in the infarct core striatum and 8% of cells in the infarcted cortex had incorporated EdU into their nuclei, significantly more than in each region in the contralateral hemisphere. In the peri-infarct striatum, 21% of all cells were EdU^+^ in the striatum and 10% of all cells were EdU^+^ in the cortex. In the contralateral striatum and cortex, just 0.5% of all cells were EdU^+^, indicating increased cell proliferation within the stroke lesion. Importantly, several EdU^+^ nuclei were co-localized with EGFP^+^ cells with pericyte-like morphology in areas where pericyte density was reduced by stroke (**Figure 6e, inserts**). Within the infarct core reductions in pericyte density in the striatum and cortex were accompanied by significant increases in the percentage of EdU^+^ pericytes compared to contralateral regions (**Figure 6g**). In peri-infarct striatum, where density was significantly reduced, we also noted a 13% increase in the percentage of EdU^+^ pericytes compared to the contralateral striatum (**Figure 6g**). We next validated these results with Ki67 staining, a standard marker for cell cycle entry and observed significant reductions in pericyte density in the infarct core and cortex, respectively (**Figure 6f, inserts**). Within the remaining populations, 12% of pericytes in the infarct striatum and 16% of pericytes in the infarct cortex were Ki67^+^, significantly more than in the contralateral hemisphere. In peri-infarct tissue, striatal and cortical pericyte density was significantly reduced and 26% of pericytes in the striatum and 10% of pericytes in the peri-infarct cortex were Ki67^+^ (**Figure 6h**). Cumulatively, these data demonstrate pericytes respond to cerebral ischemia by entering the cell cycle where pericyte density is reduced.

### Surviving ischemic pericytes drive prolonged vasoconstriction

Finally, we sought to dissect how surviving pericytes within the previously ischemic cortex regulate capillary diameter in the sub-acute phase after stroke. We first asked whether pericyte subtypes react differently to ischemia. We observed that every pericyte subtype dilated to near pre-stroke levels after 24 hours, however, three and seven days post-stroke a secondary decline in capillary lumen diameter at pericyte soma occurred within thin-strand pericyte subtypes (**Figure 7a, b**). These data suggest that while ischemic mesh pericytes recover well in the sub-acute, en passant thin-strand and junctional thin-strand pericytes remain constricted and limit blood flow long-term.

**Fig. 7.**
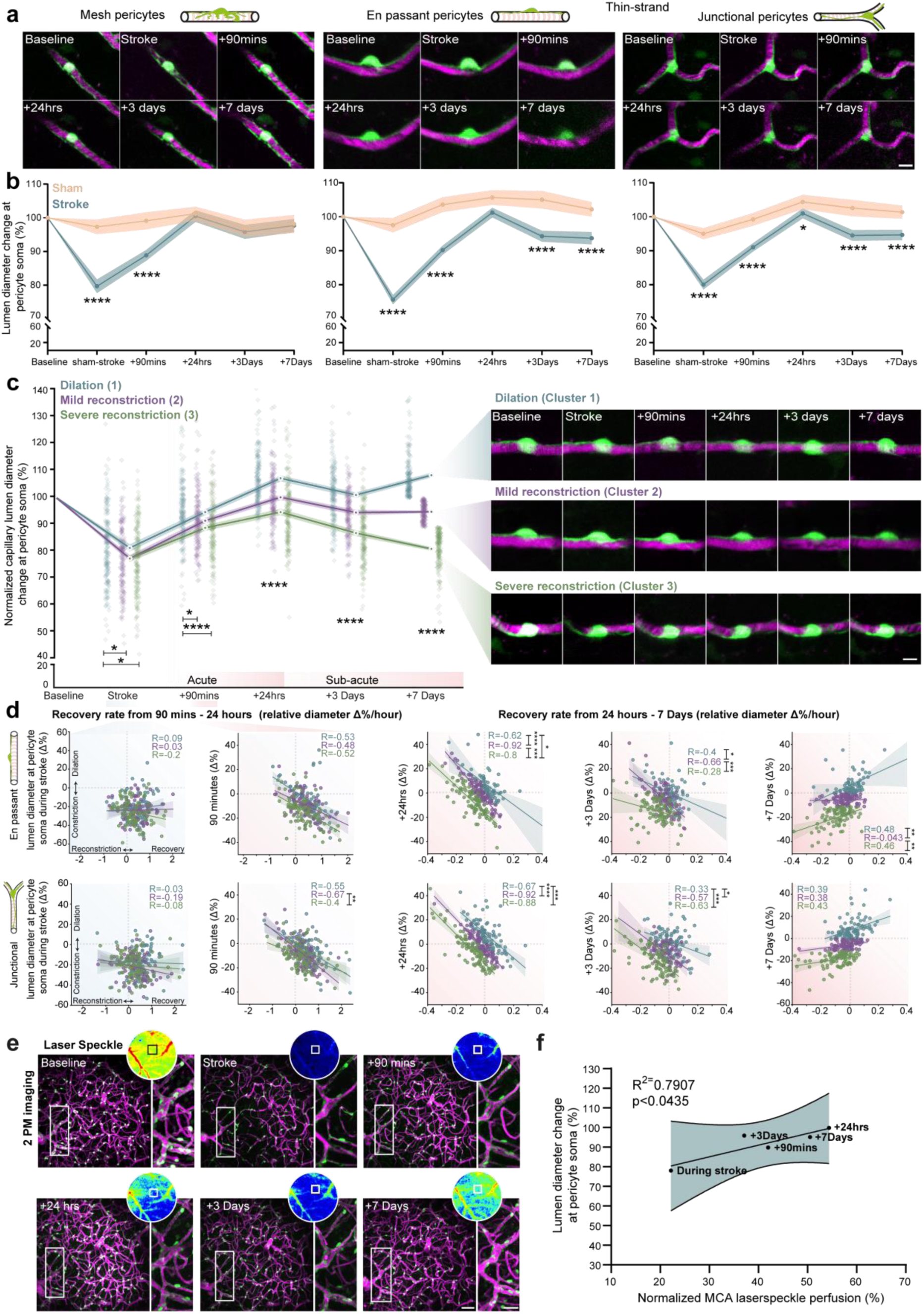
Impaired acute recovery of ischemic thin-strand pericytes drives prolonged vasoconstriction. **a)** 2-PM imaging of individual pericyte subtypes within the stroke group. **b)** Normalized lumen diameter change at pericyte soma between groups. **c)** Left, mean normalized lumen diameter change at pericyte soma within day 7 pericyte clusters which are dilating (blue), mildy reconstricting (magenta) or severely reconstricting (green) over experimental time course. Right, representative images of pericytes belonging to each cluster. **d)** Left, correlation (comparison/analysis) in between acute recovery rate and capillary diameter changes during stroke and 90mins post occlusion based on day 7 clustering. Right, correlation between subacute recovery rate and capillary diameter changes 24 hours post occlusion to 7 days post stroke based on day 7 clustering. **e)** 2-PM imaging of a single capillary bed and its position within the laser speckle field in the somatosensory cortex over experimental time course. **f)** Correlation between mean lumen diameter at pericyte soma and normalized laser speckle perfusion.. n=6 PDGFRßEGFP (stroke), n =4 (sham). Statistics, **b, c** Two-way ANOVA with Tukey’s post-hoc correction. **d**, Pearson’s rank correlation, transformed into Z-Fisher’s scores and compared using the cocor package within R. **f,** Pearson’s rank correlation. In each figure, significance codes *p < 0.05, **p < < 0.01, ***p < 0.001 and ****p < 0.0001. Scale bars **a,** 10 µm, **c,** 5 µm, **e,** 50 µm and 25 µm respectively. Data is shown as mean +/-s.e.m **b, c, d** and s.d for **f** and source data are provided as a source data file.

To disentangle how persistent constriction arises in two thirds of the thin-strand pericyte population, we grouped both thin-strand pericyte subtypes together into three equally sized clusters based on the variation of capillary lumen diameter on day 7 (Dilation, Mild reconstriction and Severe reconstriction **Figure 7c**). During stroke, thin-strand pericytes that reconstricted in the sub-acute phase (cluster 2, 3; mild and severely reconstricting populations) constricted significantly more than pericytes that were dilated in the sub-acute phase (Cluster 1). At 90 minutes post reperfusion, pericytes that dilated in the sub-acute phase constricted by 6%, significantly less than pericytes that reconstricted in the sub-acute phase. Strikingly, after 24 hours, dilating, mild reconstriction and severe reconstriction clusters of pericytes diverged significantly from one another, a pattern which strengthened on day 3 and determined day 7 post stroke clustering. These data suggested pericytes may retain an imprint of constriction during ischemia which could determine their future contractile behavior in the sub-acute phase of ischemic stroke. Moreover, mild and severe reconstricting clusters only began to diverge upon reperfusion, which suggested ischemic thin-strand pericytes might show delayed recovery of capillary diameter upon restoration of blood flow.

To explore this, we compared the constriction state at each time-point with the rate of capillary lumen diameter recovery at pericyte soma within the acute (90 mins – 24h) and subacute phase (24h – 7days) in each cluster of thin-strand pericytes. No significant correlation was found when comparing pericyte contractility levels during stroke with acute recovery in capillary lumen diameter (90 mins – 24 hours, p>0.05 for all clusters, **Figure 7d, during stroke**). After reperfusion, ischemic pericytes from all clusters recovered at a rate inversely proportional to their constriction strength in the presence of blood flow (**Figure 7d, 90 mins – 24 h**). These data strongly indicate that upon reperfusion, surviving pericytes do not maintain a permanently contracted state in the ischemic cortex.

Interestingly however, junctional thin-strand pericytes that severely reconstricted on day 7 showed a significantly weaker rate of recovery than mildly reconstricting junctional thin-strand pericytes, suggesting a distinct delayed recovery within this cluster. Moreover, we found pericytes from mild and severe reconstriction clusters in both thin-strand pericyte subtypes trended toward reconstriction significantly faster than those pericytes found in the dilating cluster on day 7 post stroke (**Figure 7d,+24 hrs and +3 Days**). Specifically, even though correlations in dilating and severely reconstricting populations of pericytes appeared similar, they trended toward opposite outcomes, with dilating clusters trending further toward dilation and constricted clusters propagating further into a constricted state. Cumulatively, these data suggest that transient ischemia chronically affects the ability of most surviving capillary pericytes to regulate the diameter of their associated capillary, with constricted pericytes dominantly limiting the dilation of their associated capillary in the sub-acute phase of ischemic stroke.

Finally, given that 57% of total vascular resistance to flow in the brain parenchyma is generated in capillaries directly adjacent to feeding arterioles, we asked whether measurements of lumen diameter variation at pericyte soma translated to an observable effect on mesoscale CBF ^24, 25^. Qualitatively, we observed that within the area covered by the cranial window, pericyte constriction and laser speckle blood flow mirrored one another closely, when pericytes were constricted, cortical blood flow was heavily reduced (**Figure 7e, Supplementary Figure 11d,e)**. We found that lumen diameter variation at pericyte soma and laser speckle blood flow were significantly correlated (**Figure 7f**) and generated an R^2^ value of 0.79, suggesting pericyte influence over vessel diameter contributes to large scale mesoscale reductions in blood flow post-stroke.

## Discussion

No-reflow of cerebral microvessels occurs in 83% of ischemic stroke survivors and results in a progressive additional loss of brain tissue ^26–30^. In the current study, we show capillary pericytes can survive transient ischemia and mediate no-reflow development by limiting reperfusion through sustained constriction of their associated capillary *in vivo*. We found pericyte constriction was widespread, most prominent at the cell soma, and occurred in both a depth and subtype specific manner during ischemia. Specifically, we observed biconcave constrictions at mesh pericyte locations and focal constrictions at thin-strand pericytes, contrasting with prior reports where focal constrictions were not found ^13^. These data may be explained by gradations of α-smooth muscle actin (αSMA) expression from low to higher order capillary pericytes and that thin-strand pericytes on higher order capillaries still express a small pool of αSMA near the cell soma^7^.

Despite this, optogenetic stimulation of pericytes lacking αSMA still produces decreases in blood flow, suggesting pericyte contractility may be mediated by an array of contractile proteins and does not solely rely on αSMA expression ^5^. We observed 87% of ischemic pericytes constricted *in vivo*, this near ubiquitous constriction is consistent with ex vivo models of chemical ischemia that report pronounced constriction of pericytes ^12^, and could be mediated by amplified Ca^2+^ influx and vasoactive agonists such as endothelin-1 and 20-HETE ^15, 19, 31^. While experiments performed in ischemic brain slices show pericytes may constrict the vessel lumen by up to 80%, our findings align with retinal ischemia models and suggest maximal pericyte constriction is around 60% of baseline ^6^. This proved sufficient to significantly impede and completely arrest blood flow *in vivo* within 24 hours after ischemic stroke and could explain ‘nodal-like’ constrictions observed in histological brain tissue ^12^. While we cannot rule out the effect a loss of upstream flow has on vessel diameter during stroke, most pericytes remained constricted after reperfusion, strongly arguing against a loss in upstream flow mediating the effect we observe at pericyte soma.

Concomitantly, we observed blebbing and cytoplasmic shedding of pericytes within the ischemic territory, suggesting they are structurally impaired. These data suggest both the intrinsic cellular function and the signal relay function of the pericyte network could be critically compromised shortly after ischemia ^9^. To test for potential membrane damage, we examined nuclear EGFP intensity in pericytes because EGFP can freely translocate through nuclear pores^32^. We reasoned that EGFP should evenly distribute in pericyte nuclei with a uniform volume, but may leak from pericytes with compromised cell membranes. We observed less EGFP in damaged pericytes, suggesting cellular cytoplasmic leakage after pericyte membrane damage is an acute reaction to ischemia. These data mirror prior publications displaying bleb formation in pericytes and a loss of membrane integrity following anoxic depolarization ^16, 18, 33^. Importantly however, we observed individual pericytes with ischemia associated blebbing remaining visible using longitudinal imaging, implying pericytes may possess the capacity to recover from ischemic membrane damage, as described for other cell types^10, 16, 33, 34^.

Pericyte susceptibility to ischemia has recently been investigated extensively in brain slices leading to the hypothesis that pericytes may irreversibly contract around capillaries and die rapidly ‘in-rigor’ ^16^. This would preclude stroke therapy aimed at ameliorating no-reflow and is therefore of critical clinical importance. Consistent with prior reports, we observed the maxima of pericyte death to occur acutely. However, our longitudinal *in vivo* findings argue against the majority of pericytes being in a death-induced contractile state in the ischemic cortex. Principally, while most pericytes in our dataset survived, their persistent constriction after reperfusion may instead indicate chronic cellular dysfunction, which suggests therapeutically targeting pericytes could ameliorate no-reflow after ischemic stroke ^16, 17^.

Pericyte survival may rely on their cellular plasticity and ability to metabolize limited perivascular glycogen stores found in astrocytic end-feet, and by harnessing glycolytic metabolic pathways to endure sustained oligemia after reperfusion ^6, 35–37^. While more research is required to decipher the intracellular mechanisms underpinning pericyte survival, region dependent pericyte survival may be explained by the presence of lepto-meningeal collaterals (LMCs) which are found in 80% of stroke patients and provide blood flow from anterior communicating and posterior communicating arteries quickly after blockage of the MCA in mice ^38, 39^ ^40^.

Though we demonstrate that pericytes can survive transient cerebral ischemia, they may be unable to effectively regulate capillary diameter to ensure adequate tissue perfusion, as we demonstrate two thirds of capillary pericytes carry a lasting imprint of constriction that negatively affects blood flow long-term. Recent insights depicting pericytes as metabolic sentinels may explain pronounced bi-phasic thin-strand pericyte contraction and indicate why capillary stalls occur most frequently in upper layers of the cortical microvasculature after ischemia. In the absence of an activity induced metabolic milieu released from neurons and astrocytes, deep capillary pericytes may fail to transduce retrograde dilatory signals to arterioles, effectively creating a more contractile basal tone within the capillary bed ^41–43^. Thus, through both constriction and impaired signaling, structurally impaired ischemic pericytes may transmit resistance to blood flow, creating a potent environment for blood flow stagnation, neutrophil obstruction and increased leukocyte-endothelial interactions^44, 45^.

Thin-strand pericytes constricted most heavily within our dataset. Moreover, thin-strand pericytes were associated with most of the capillary stalls and were responsible for chronic reconstriction of capillaries within the first week of stroke. These findings highlight that thin-strand pericytes in particular, may be profoundly susceptible to ischemia.

Finally, we observed that reductions in pericyte density elicited local proliferation of pericytes in the sub-acute phase. This data is entirely consistent with recent publications describing an evolutionarily conserved wound healing proliferative profile of activated pericytes ^46, 47^. Thus, a significant proportion of ischemic pericytes may be coerced into wound-healing, angiogenic, pro-regenerative transcriptomic profiles by their surrounding inflammatory milieu and lack of pericyte-pericyte contact, which may further hinder their ability to effectively regulate blood flow.

In summary, the current study offers a critical, long-needed *in vivo* insight into the role pericytes play in no reflow after ischemic stroke. Pericytes are particularly resistant to cerebral ischemia, but constrict cerebral capillaries during and after transient MCA occlusion and contribute to no-reflow development, principally within the first 24 hours post-stroke. Within the first few days after cerebral ischemia, pericytes proliferate and regenerate the capillary pericyte network, but continue to restrict blood flow to the ischemic brain long-term. Our data suggest structural and functional impairment, not immediate death of cortical pericytes mediates no-reflow after transient cerebral ischemia. Therefore, pericytes may represent a novel therapeutic target for the treatment of no-reflow after ischemic stroke.

## Methods

### Animal handling

Adult male *PDGFRß*-EGFP^ret/ret^*^WT^* mice alone, or crossed with (Tg.(*Cspg4*-DsRed) 1Akik/J mice and male C57BL/6J mice (Jackson Laboratory) between 6-22 weeks old were used for this study. The animals were housed under a 12 h/12 h light/dark cycle and were provided with pellet food and water ad libitum. All experiments were approved by the Ethics Review Board of the Government of Upper Bavaria (protocol number 81-017).

### Chronic cranial window implantation

Animals were anesthetized with Medetomidine 0.5 mg/kg, Midazolam 5 mg/kg and Fentanyl 0.05 mg/kg (MMF) by i.p. injection and a craniotomy was performed over the right parietal cortex. The cranial bone flap and exposed dura mater were carefully removed and a round, 4 mm cover glass was fixed to the skull using Vetbond^®^. A custom titanium ring was attached to the skull using UV hardening dental cement ^48^. Anesthesia was antagonized with Atipamezol 0.5 mg/ml, Naloxon 3 mg/ml, Flumazenil 5 mg/ml (ANF) i.p. and mice were then placed in a heating chamber at 32°C for 2 hours until fully recovered.

### Transient filament middle cerebral artery occlusion

30 minutes prior to surgery mice were injected intraperitoneally (i.p) with 0.1 mg/kg buprenorphine to act as pre-operative analgesia. The middle cerebral artery (MCA) was occluded with an intraluminal filament for 60 minutes as previously described ^49^. MCA occlusion (MCAo) was verified in each animal by intraoperative monitoring of cerebral blood flow (CBF) by either laser speckle contrast imaging (for In vivo experiments, LSCI; see below) or laser Doppler fluxmetry (for *ex vivo* experiments) and subjected to examination of neurological deficits using the Bederson score. A previously used post-operative care regimen was modified to maximize and preserve the survival of animals following MCAo ^50^.

### In vivo imaging of pericytes and blood-flow after stroke

NG2DsRed^+/-^PDGFRßEGFP^+^ mice were injected subcutaneously with 100 mg/kg of medetomidine and 50 µl of 25 mg/ml (3000 MW) TexasRed^®^ neutral anionic dextrans. Fifteen minutes later low levels (0.5%-0.75%) of isoflurane in 1:1 oxygen and N_2_ were used to maintain a light anesthesia.

Baseline cortical CBF recordings were assessed using LSCI (Pericam PSI, Sweden). Mice were then transferred to a Zeiss 7MP microscope (Carl Zeiss AG, Germany), equipped with a Chameleon Ultra Ti:Sapphire laser (Coherent^®^, USA), and a 20×1.0 NA water immersion objective. Z-stacks of regions of interest (ROIs) with dimensions of 425 µm x 425 µm were acquired from the brain surface down to a depth of 300 µm. ROIs were chosen based on minimal pial vessel coverage that highlighted capillary beds within the cortex and coordinates for each ROI were determined based on drilled indentation landmarks, which were placed on the cover glass prior to cranial window implantation (**Supplementary Figure 1**).

After 3-4 days, mice were subjected to either stroke or sham surgery. Stroke induction was confirmed if LSCI recordings showed a consistent, >70% reduction in CBF. Within the area of confirmed reduced perfusion, two ROIs were selected for *in vivo* 2-photon microscopy. At the end of the experiment, mice were injected with MMF i.p. and sacrificed. Brains were harvested for immunostaining.

### Longitudinal vessel lumen diameter and pericyte analysis by in vivo 2-photon imaging

To analyze cortical pericytes and the associated microvasculature image sets were separated into 4 x 50 µm stacks of increasing depth. Pericytes were divided into the following sub-types: thin-strand pericytes, junctional pericytes, and mesh pericytes based on previously published criteria ^20^. The diameter of pericyte associated vessel lumens were measured in FIJI (ImageJ v1.53f51) at the left, middle, and right side of the cell soma in individual z-sections. Comprehensive analysis of subtype-specific vessel diameter changes was executed by a custom written script in MATLAB environment (2021b, MathWorks, MA, USA).

### Capillary stall analysis by in vivo 2PM imaging

Capillary stalls were identified by shadows in the vessel lumen not stained by TexasRed^®^ according to the following criteria: If the shadow in the capillary (mainly caused by red blood cells; RBC) was present for 3 or more consecutive frames of scanning (4.8 seconds) the capillary was deemed ‘stalled’. This is because the average flux of 50 RBC/s is reported in pia 0-100 µm below the surface ^24^. Stalls were termed “pericyte associated” if they appeared at a distance less than 5 µm from the pericyte soma (**Supplementary Figure 5**).

### Perfusion of mice and tissue collection

Animals were anesthetized with MMF and transcardially perfused with PBS followed by 4% PFA until the liver was devoid of blood. Brains were extracted, post-fixed in 4% PFA over night and then stored in 1XPBS at 4°C until further use. Brains were mounted in 4% agarose and sectioned using a vibratome (Leica VT1200S, Leica Microsystems, Germany) to create 100 µm thick brain slices. For fresh frozen sections, brains were frozen on dry ice and serially sectioned in TissueTek using a cryostat (CryoStar NX70, Thermofisher, USA) into 20 µm brain slices mounted on microscope slides and stored at-20°C until further use.

### Analysis of pericyte damage after transient cerebral ischemia

Three PFA-fixed brain sections from PDGFRßEGFP mice (n=4) subjected to fMCAo containing the MCA territory were incubated in a blocking solution and permeabilizing primary antibody buffer solution (1% BSA, 0.1% fish-skin gelatin, 0.1% Triton X-100, 0.05% Tween 20 in 1XPBS) with 1:100 dilutions of goat anti-collagen IV (ab235296, Abcam), rabbit anti-Aquaporin IV (AB2218, Merck Millipore), GFAP-Cy3 (C2905, Sigma), or iba1 (019-19741, WAKO) antibodies on a rotary shaker at 4°C for 2-3 days. Sections were then washed in 1XPBS 3X for 30 minutes and incubated with a secondary antibody buffer mix (in 2% BSA, 2% FCS, 0.2% fish-skin gelatin in 1XPBS) containing a donkey anti-rabbit AlexaFluor^®^ 594 (Jackson) and donkey anti-goat AlexaFluor^®^ 647 (Jackson) (1:300) antibody for at 4°C on a rotary shaker two days. Sections were then washed three times in 1XPBS and during the last washing step DAPI was added at a concentration of 1:1000 for 30 minutes to stain the nuclei. Washing using 1XPBS was repeated three times for 30 minutes prior to mounting the sections on glass coverslips with EverBrite™ mounting medium. Using high-resolution confocal imaging (LSM 880, Carl Zeiss, Germany) with a 100x oil objective (Epiplan-Neofluar 100x/1.3 Oil Pol M27), pericytes in the infarct core region and contralateral hemisphere were imaged for downstream analysis. Pericytes were separated by sub-type definition (thin-strand, mesh, junctional) and reconstructed using IMARIS^®^ software (Oxford Instruments, Bitplane, Switzerland). Surface creation of DAPI^+^ nuclei allowed measurement of nucleus volume and sphericity.

### Analysis of pericyte death, coverage and proliferation bodies

Twenty-five C57BL/6J mice allocated to three experimental cohorts (fMCAo, sham surgery or no surgery) in a randomized and blinded manner were sacrificed on day 1 or day 3 after surgery (n=5 per group). After serial cryostat sectioning, brain slices were rehydrated, fixed in 4% paraformaldehyde, and blocked and permeabilized in 2% BSA, 2% FCS, 0.2% fish-skin gelatin in 1XPBS and 0.03% Triton x-100 for 1 hour at room temperature. Slices were incubated with primary antibody buffer mix consisting of gtPDGFRß (1:100; AF1042, R&D Systems), rbCollagen IV (1:250, Ab19808, Abcam) over night at 4^°^C. On the next day brain sections were washed three times with 1XPBS and incubated with donkey anti-goat Alexafluor^®^ 488 (1:1000) and donkey anti-rabbit Alexafluor^®^ 647 (1:300) secondary antibodies. Sections were then stained with the ApopTag^®^ Red In Situ Apoptosis Detection kit following manufacturers guidelines. During the secondary incubation of the dioxygenin conjugate, sections were incubated with DAPI to stain nuclei. Brain slices were mounted on glass slides and five regions of interest were imaged by confocal microscopy (Zeiss LSM 880, Oberkochen, Germany) in the infarct, peri-infarct and contralateral areas in two brain slices per mouse using a 40x oil objective (EC Plan-Neofluar 40x/1.30 Oil DIC M27, Zeiss).

### Bregma dependent analysis of pericyte survival 3 days post-stroke

The brains of four male PDGFRßEGFP^+^ mice, subjected to a one-hour transient fMCAO, were removed and prepared for immunostaining of PFA fixed tissue. Three brain sections per mouse ranging from the posterior MCA territory (bregma-0.9/-1.2 mm), MCA territory (bregma 0/0.1 mm) to anterior cortex (bregma +2.5/2.71 mm) were immunostained. Sections were incubated with primary antibody buffer solution with 1:200 rabbit anti-NeuN (ab177487, Abcam), goat anti-Aquaporin IV (AB2216, Merck Millipore), GFAP-Cy3 (C2905, Sigma), or iba1 (019-19741, WAKO) for 2 days on a rotary shaker at 4°C. Sections were washed three times in 1XPBS and incubated with a secondary antibody mix (see previous description) in 1XPBS and 1:300 donkey anti-rabbit AlexaFluor^®^ 647 on a rotary shaker at 4°C for 2 days. Sections were then washed with 1XPBS, incubated with DAPI and mounted as previously described.

Sections were imaged with a confocal microscope (Zeiss LSM 880, Oberkochen, Germany) by using a 5X air objective (EC Plan-Neofluar 5x/0.16 Pol M27, Zeiss). For analysis of pericyte population survival, images were imported into IMARIS^®^ and the green and the red channel were separated. Using the red NeuN^+/-^ signal within the tissue, three areas were traced: Infarct core, ipsilateral hemisphere and contralateral hemisphere. The green PDGFRßEGFP channel was subjected to 3D surface creation of spots representing cells in IMARIS^®^ based on a minimum feature size of 8 µm which allowed calculation of mural cell number in each distinct region: Infarct core, ipsilateral hemisphere and contralateral hemisphere. Finally, cell spots were deleted based on their vascular location (arteries, veins) or doublets; leaving a count of capillary level PDGFRßEGFP^+^ pericytes.

### Cell cycle entry analysis

Four male PDGFRßEGFP^+^ mice, subjected to a one-hour transient fMCAo were sacrificed three days after stroke, brains were removed and prepared for immunostaining of PFA fixed tissue. Two brain sections including the MCA were chosen for further immunostaining. Briefly, sections were incubated with primary antibody buffer with 1:200 rabbit anti-Ki67 (D3B5, Cell Signaling) for 2 days on a rotary shaker at 4°C. Sections were washed three times in 1XPBS and incubated with a secondary antibody mix in 1XPBS and 1:300 donkey anti-rabbit AlexaFluor^®^ 647 on a rotary shaker at 4°C for 2 days. For confocal analysis, three regions of interest were chosen in the striatum and cortex in the infarct core, the peri-infarct region and corresponding contralateral brain regions. The pericyte density was assessed by quantifying the number of PDGFRßEGFP^+^ cells at the capillary level.

### 5-Ethynyl-2’-deoxyuridine (EdU) administration and staining

Four PDGFRßEGFP^+^ male mice were subjected to one-hour transient fMCAo. 24 hours post stroke, each animal was injected intra-peritoneally every 8 hours with 80 mg/kg (2 mg EdU/injection) EdU dissolved in *aqua ad injectabilia* (Chemie, Berlin) until sacrifice at 72 hours. Brain sections corresponding to the posterior MCA territory, the MCA territory and anterior cortex were stained with Click-iT ™ Plus EdU Cell proliferation kit (C10640, Thermo Fisher Scientific) following manufacturer’s instructions and mounted using EverBrite^TM^ mounting medium as previously described. Using the 40X oil objective at the confocal microscope, three volumetric images per brain section and region were acquired. For image analysis total number of pericytes, EdU^+^ cells and DAPI^+^ cells were counted per region of interest. Finally, the number of total pericytes per ROI was normalized to the contralateral pericyte number to display pericyte density in relation to the contralateral hemisphere and the total number of EdU^+^ pericytes were displayed as a percentage of the remaining pericyte density within each region (infarct core/periinfarct striatum, cortex). Δ

### FACS isolation of pericytes and bulk RNAseq analysis

After three days of either stroke or sham surgery mice were sacrificed (in pairs, one stroke and one sham) via cervical dislocation, brains were extracted and placed on ice cold 2% FBS in 1XPBS solution in a petri-dish. In a brain mold, distinct regions corresponding to the infarct core, ipsilateral hemisphere (of stroke and sham animals) and contralateral hemisphere were mechanically and enzymatically dissociated to release living EGFP^+^ and DsRed^+^ cell populations from the microvasculature using a modified version of a previously published protocol ^21^. DAPI^-^ EGFP^+^ DsRed^+^ cells were sorted and collected from each region and collected into pools of 50 in a 96 well plate using a Sony SH800 cell sorter (Sony, Japan). For detailed information regarding the FACS isolation and gating strategy, see supplementary methods and supplementary figure10.

Methodology for bulk sequencing library preparation was performed as described in a prior publication ^51^. Briefly, pools of 50 DsRed/EGFP^+^ cells isolated from the brains of day 3 stroke or sham animals were sorted into 96 well plates, already filled with 4 μL lysis buffer containing 0.05% Triton X-100 (Sigma), ERCC (External RNA Controls Consortium) RNA spike-in Mix (Ambion, Life Technologies) (1:24000000 dilution), 2.5 μM oligo-dT, 2.5 mM dNTP and 2 U/μL of recombinant RNase inhibitor (Clontech) then spun down and frozen at-80°C. For detailed information of library preparation using the SmartSeq2 pipeline, Seurat cell cycle and gene ontology analysis see supplementary methods.

### Statistical analyses

Unless otherwise stated, all datasets are presented as mean ± SD and were assessed for normality using the Shapiro-Wilk normality test and analyzed statistically using PRISM version 9.3.1. For all longitudinal assessments and *in vivo* experimentation, Two-way ANOVA using the Greenhouse-Geisser correction method were used and subjected to Tukey’s multiple comparisons test. All pairwise comparisons were performed using either a two-tailed Student’s *t*-test or Welch’s *t*-test. For non-normally distributed and independently grouped data, the Kruskall-Wallis test was used and the data presented as median ± interquartile range. For comparisons of Pearson’s Rank r values generated within the rstats package within python, the data were transformed using a Z-fisher transformation and compared using the cocor package within Rstudio v4.1.2 (Rstudio team, USA). For all statistical analyses a *p* value < 0.05 was used as the significance threshold.

## Supporting information

Source data for all figures

## Author contributions

All authors have read and approved the final submission. **J.S:** Conceptualization, experimental design, collection of the data, interpretation and analysis of the data, drafting the article and critical revisions to the article. **N.P:** concept and study design, critical intellectual input regarding the article and corrections to the draft and final manuscript submission. **S.F:** implantation of chronic cranial windows, assistance in figure creation and interpretation of the data. **D.V:** analysis of critical *in vivo* data. **U.M:** filament middle artery occlusion of the *in vivo* experimental cohorts and FACS cohorts, injection of EdU in pericyte replication studies. **S.BG:** Processing and data evaluation of BulkSeq RNA data, transcriptomic profiling, string database analyses and gene ontology analysis and critical input of the sequencing data during the figure creation process. **B.B:** processing of the brain tissue for FACS isolation of pericytes and establishment of the FACS gating strategy. **B.S, B.G**: Critical input for analyses of laser speckle contrast imaging. **F.L, P.B:** analysis of pericytes within 2-photon dataset. **I.K, A.W**: MRI measurement and representation of infarct volumetry within the in vivo experimental cohort. **O.G:** Establishment of the FACS gating strategy for the isolation of pericytes and editing and critical input of the final manuscript. **A.L:** Editing and contribution to final manuscript submission.

## Acknowledgments

This work was supported by the Deutsche Forschungsgemeinschaft (DFG, German Research Foundation, project number: 457586042). For the sorting studies, we are grateful for support from the “Flow Cytometric Cell Sorter Sony SH800 Core Unit” run by the Department of Vascular Biology at the Institute for Stroke and Dementia Research (ISD) and SyNergy EXC2145.

---

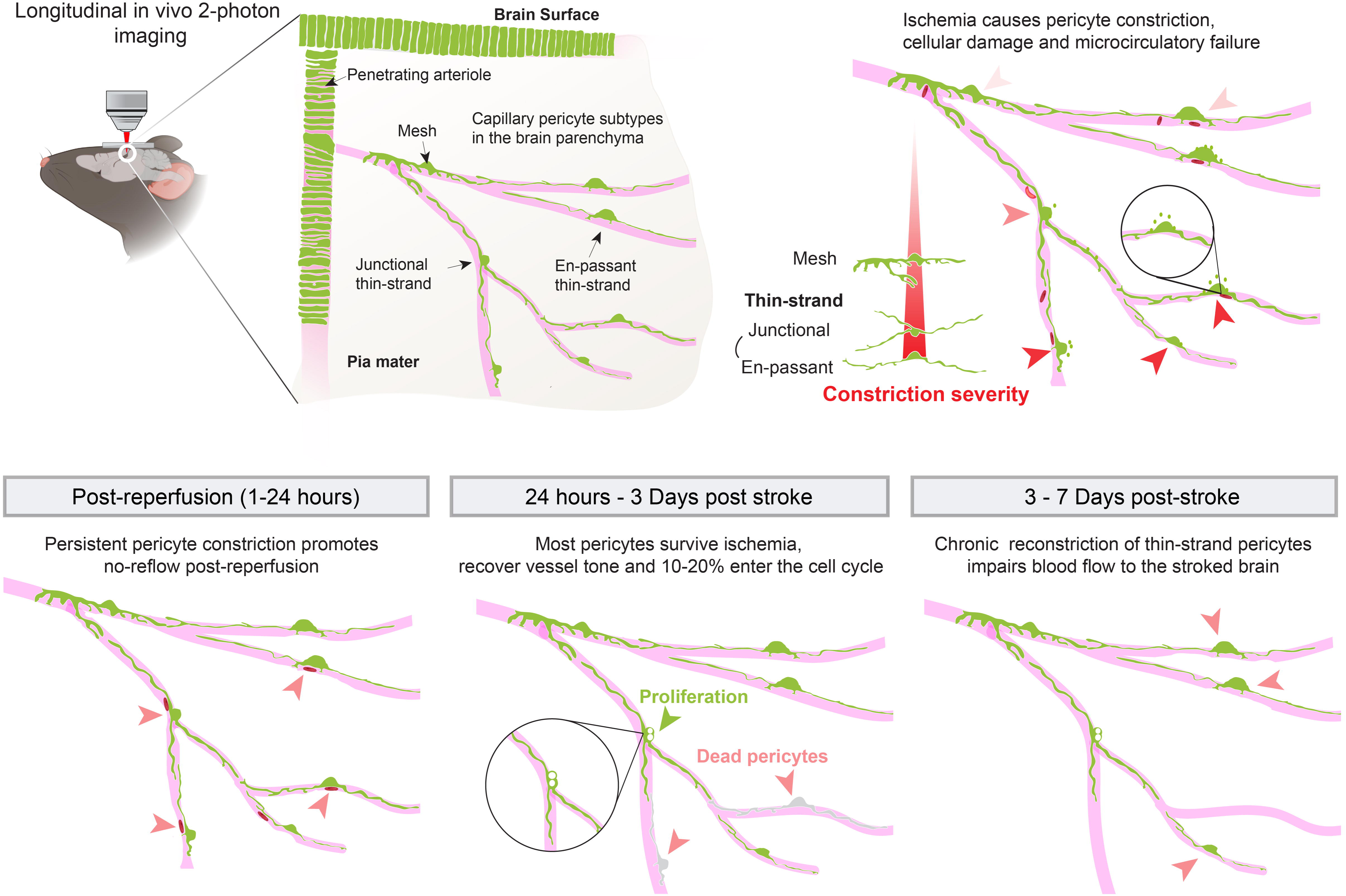

