## Supplementary figures and methods for "Continued dysfunction of capillary pericytes promotes no-reflow after experimental stroke *in vivo*": 060323_Supplementary Figures and legends_For BioRchiv.docx

**Supplementary Figures and text**

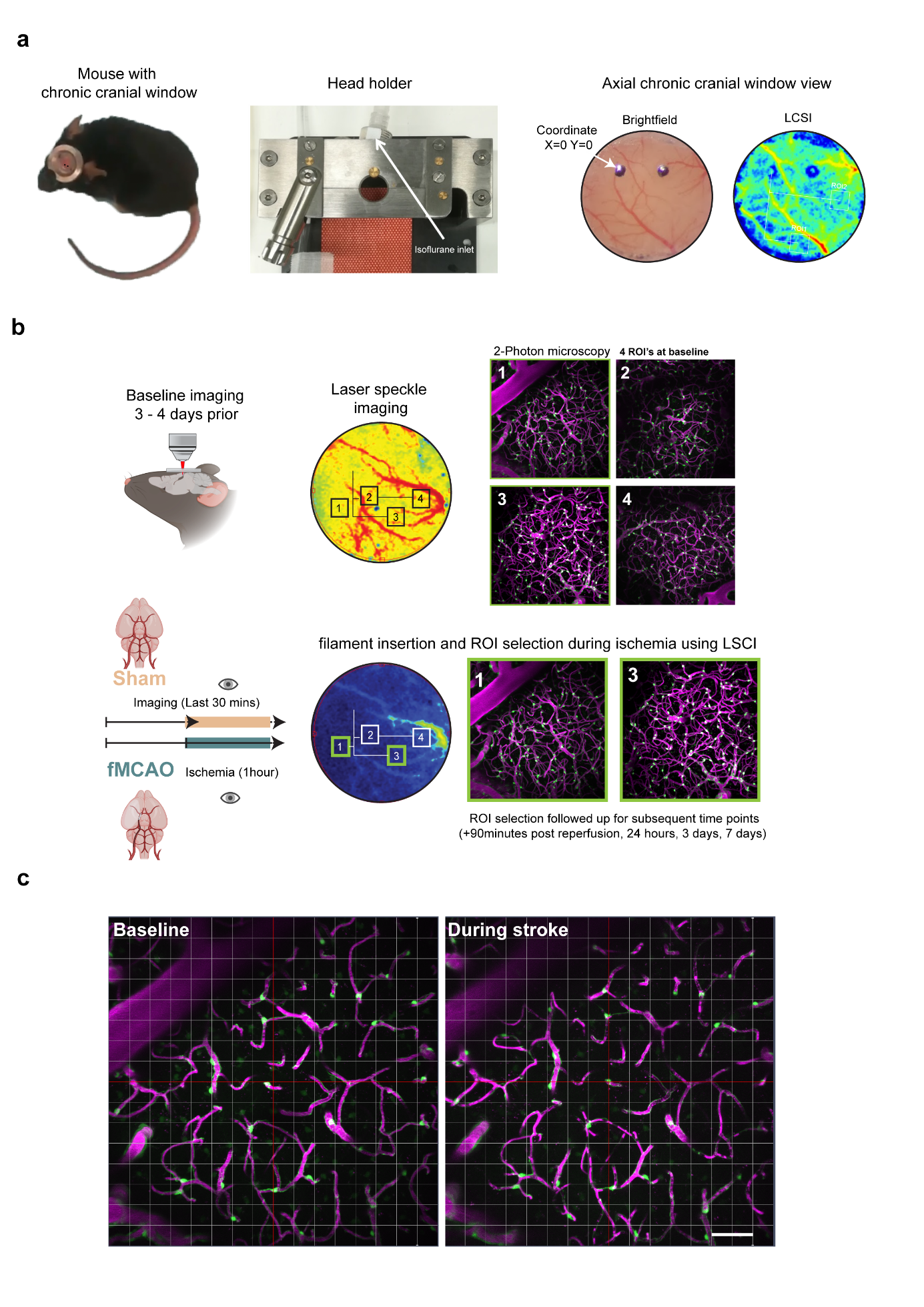

#### Supplementary Figure 1 Longitudinal imaging methodology and pixel matching for pericyte localization.

**a)** PDGFRßEGFP^+^ mouse with chronic cranial implanted, head holder and custom drilled navigation dots filled with permanent ink, with the left dot serving as coordinate 0 for bright field, LSCI and 2-photon imaging. **b)** Region of interest acquisition (ROI) and selection following during ischemia, selected of ROIs based on their perfusion by the distal MCA branch over the somatosensory cortex. **c)** Pixel matching in X-Y using a 1024x1024 line grid used across all time-points. Scale bar 50 µm.

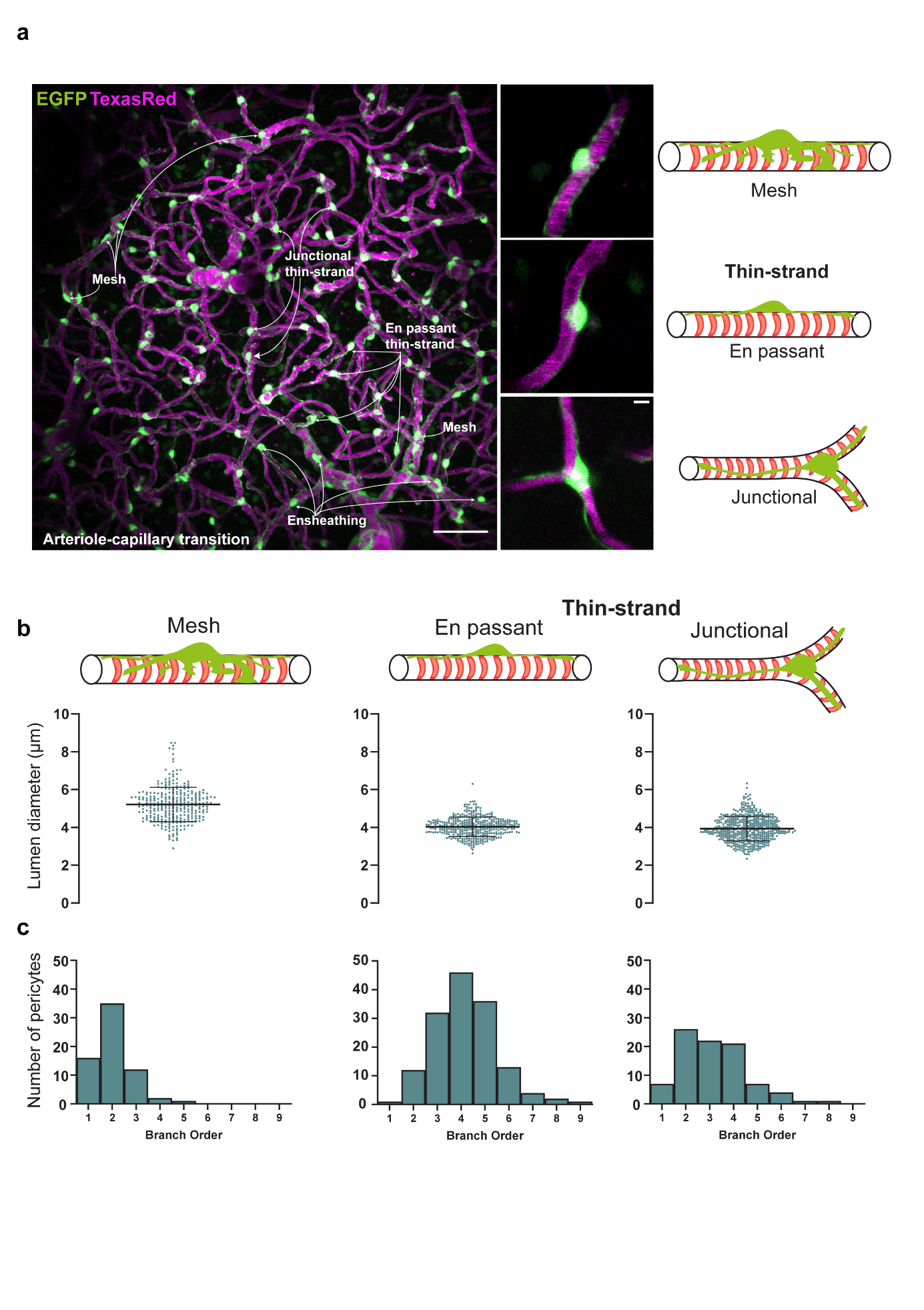

#### Supplementary Figure 2 *In vivo* characterization of capillary pericytes at baseline.

**a)** 2-PM maximum intensity projected z-stack of a typical capillary bed perfused by a penetrating arteriole used for *in vivo* analyses. White text identifies pericyte subtypes emanating from the penetrating arteriole: Ensheathing, Mesh, En passant and Junctional thin-strand pericytes. **b)** Upper, Schematic of morphological features of pericyte subtypes analyzed. Lower, Mean average vessel lumen diameter measured at each pericyte subtype soma. **c)** Branch order histograms of each pericyte subtype, n=11 PDGFRßEGFP mice. Scale bars, **a,** 50 µm and 5 µm. Data is shown as mean +/- s.d.

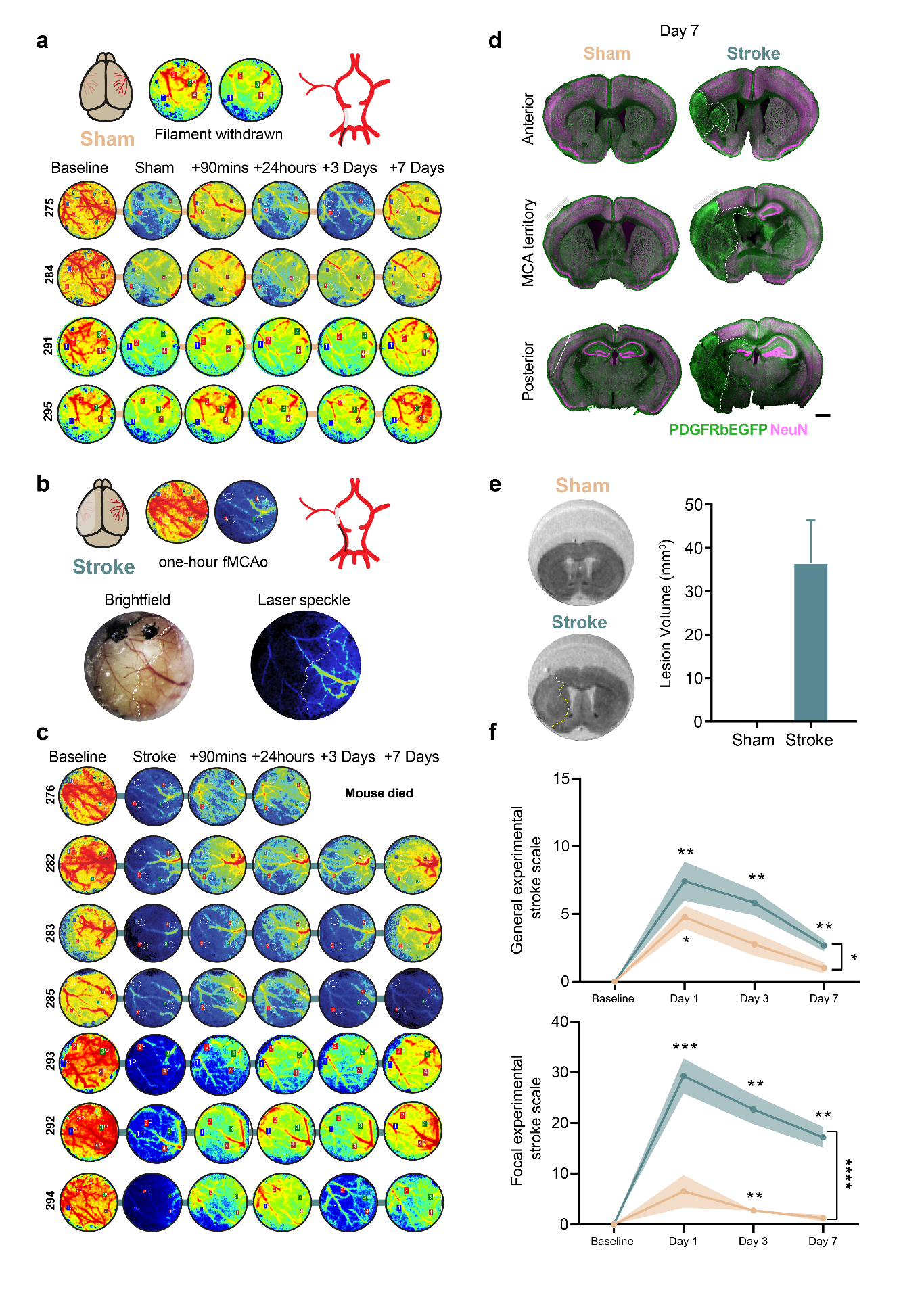

#### Supplementary Figure 3 Blood flow reduction, ischemic infarction and neurological deficits within longitudinal imaging cohort.

**a)** Raw laser speckle fluxmetry in sham group. High and moderate levels of flow represented in red and yellow respectively, while low flow is represented in green and blue. **b)** Bright-field microscopy view of the cranial window of an animal during stroke, perfused areas are a pink red color, while non-perfused tissue is white (upper). **c)** Laser speckle flow during fMCAo, white line indicates the MCA territory occluded by the filament (left). **d)** Caudal-Rostral axis immunostaining of neurons (NeuN, magenta) and PDGFRßEGFP (green) in sham and stroke animals respectively, gray bars indicate location of cranial window. **e)** Lesion volume on day 7 post-stroke/sham measured in fixed brains using MR imaging. **f)** Experimental stroke scale behavior tests throughout experimental timecourse. In each figure, significance codes *p < 0.05, **p < < 0.01, ***p < 0.001 and ****p < 0.0001. n=7 PDGFRßEGFP stroke mice, 4 sham mice, Scale bars, 1 mm. Data is shown as mean ± s.d.

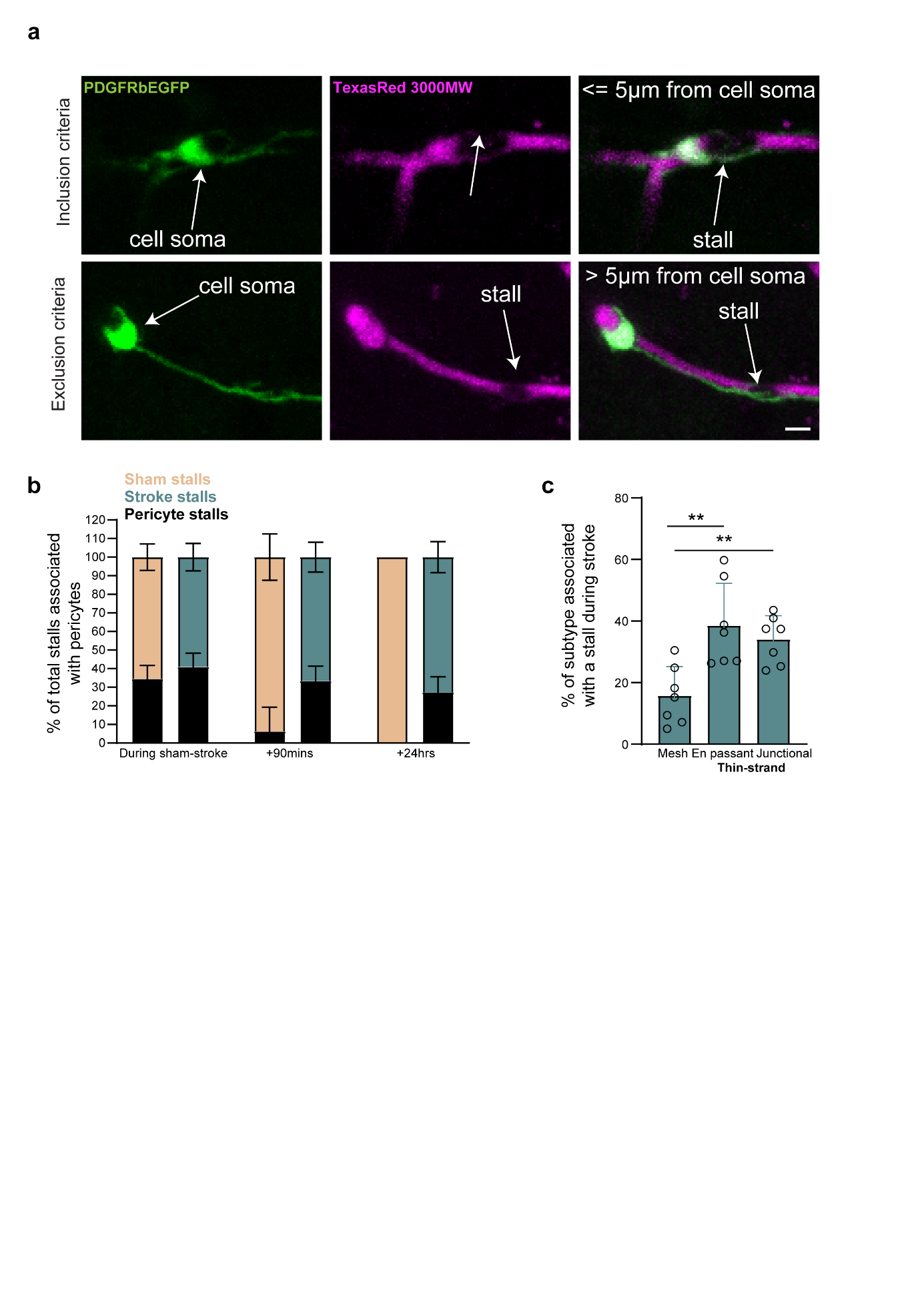

#### Supplementary Figure 4 Capillary pericytes represent a third of all capillary stalls in the ischemic brain during stroke and post-reperfusion.

**a)** Inclusion and exclusion criteria for association of capillary stalls with pericytes. **b)** Percentage of pericyte stalls as a fraction of total stall number during either stroke or sham surgery up to 24 hours post surgery in both groups. **c)** Percentage of each pericyte subtype associated with a stall during stroke.. n=7 stroke mice and 4 sham mice ~1400 pericytes stroke-890 sham-510 across 6 timepoints. Scale bars, 5 µm. Data is shown as mean +/- s.d.

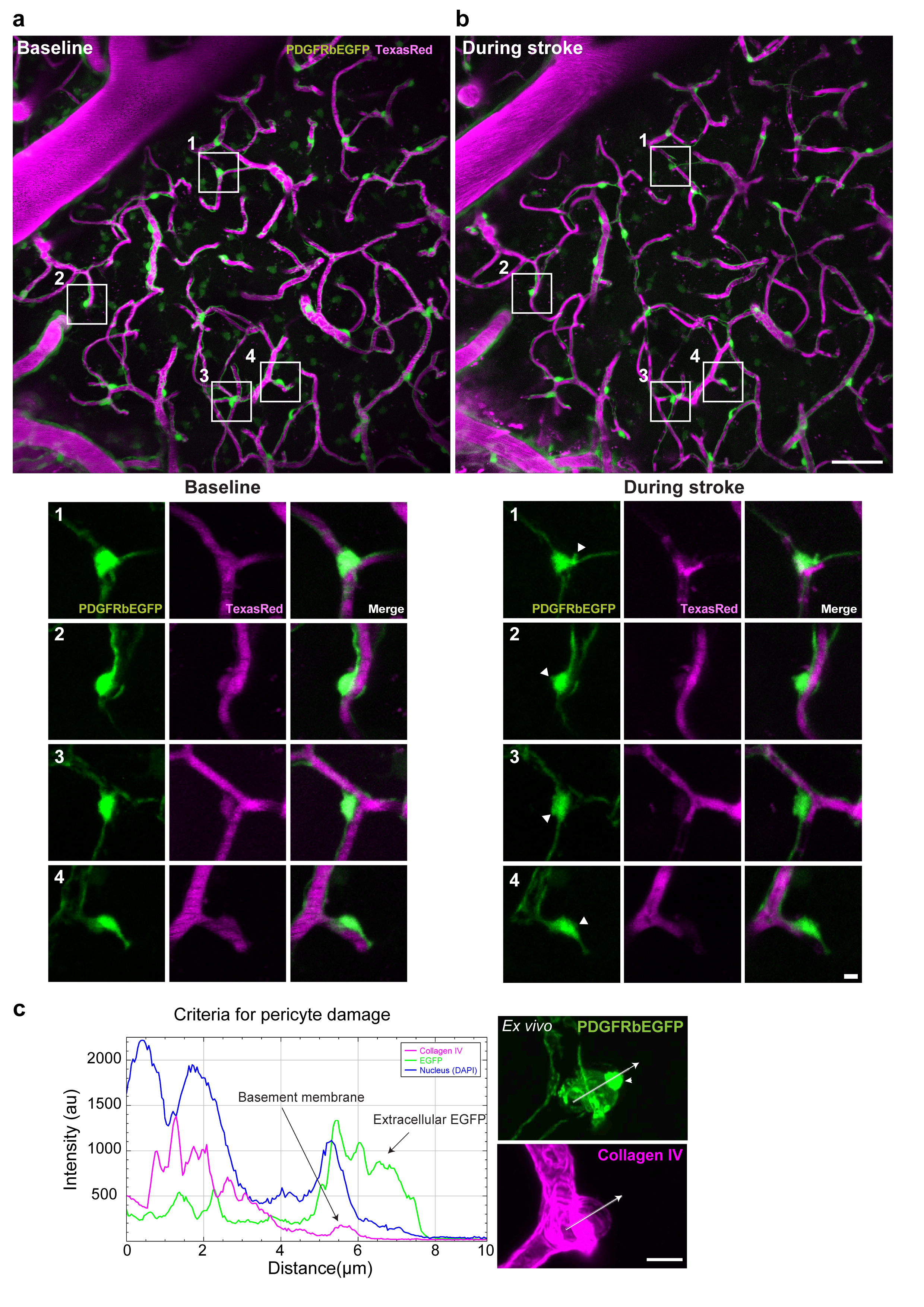

#### Supplementary Figure 5 Ischemia induces bleb formation in pericytes *in vivo*.

**a)** Upper, maximum intensity projection of a capillary bed imaged at baseline with individual pericytes highlighted in white boxes. Lower, magnification of pericyte morphology at baseline (EGFP) and vascular flow (Magenta) of pericytes 1-4 **b)** Upper, field of view imaged in A during ischemia. Lower, magnification of pericyte morphology while pericytes experience ischemia, white arrows indicate formation of blebs that were not present during baseline imaging in pericytes 1-4. **c)** Line profile measurement of EGFP leakage beyond the collagen IV^+^ basement membrane as criteria for pericyte damage. Scale bars, **a,b,** 50 µm, 5 µm **c,** 10 µm.

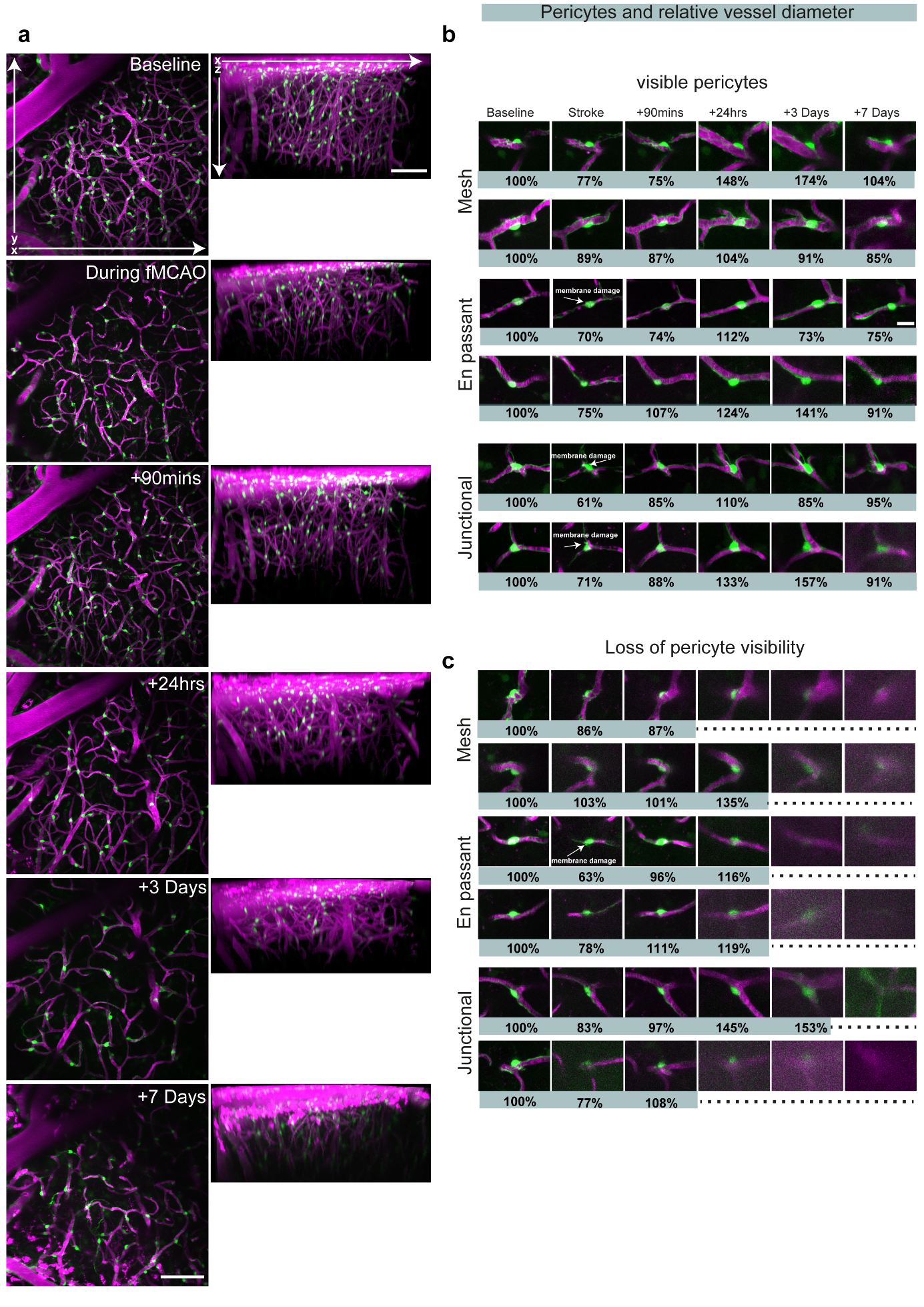

#### Supplementary Figure 6 Depth dependent loss of pericyte visibility in the ischemic cortex.

**a)** XY and XZ 2-PM maximum intensity projections of the capillary bed imaged in a stroked PDGFRßEGFP mouse over the experimental timeline. **b)** Surviving pericytes and their relative diameters over time within the capillary bed imaged. **c)** Degenerating pericytes and respective vessel diameters within the capillary bed imaged. n= Animal 294, PDGFRßEGFP stroked mouse. Scale bars, 50 µm, 10 µm.

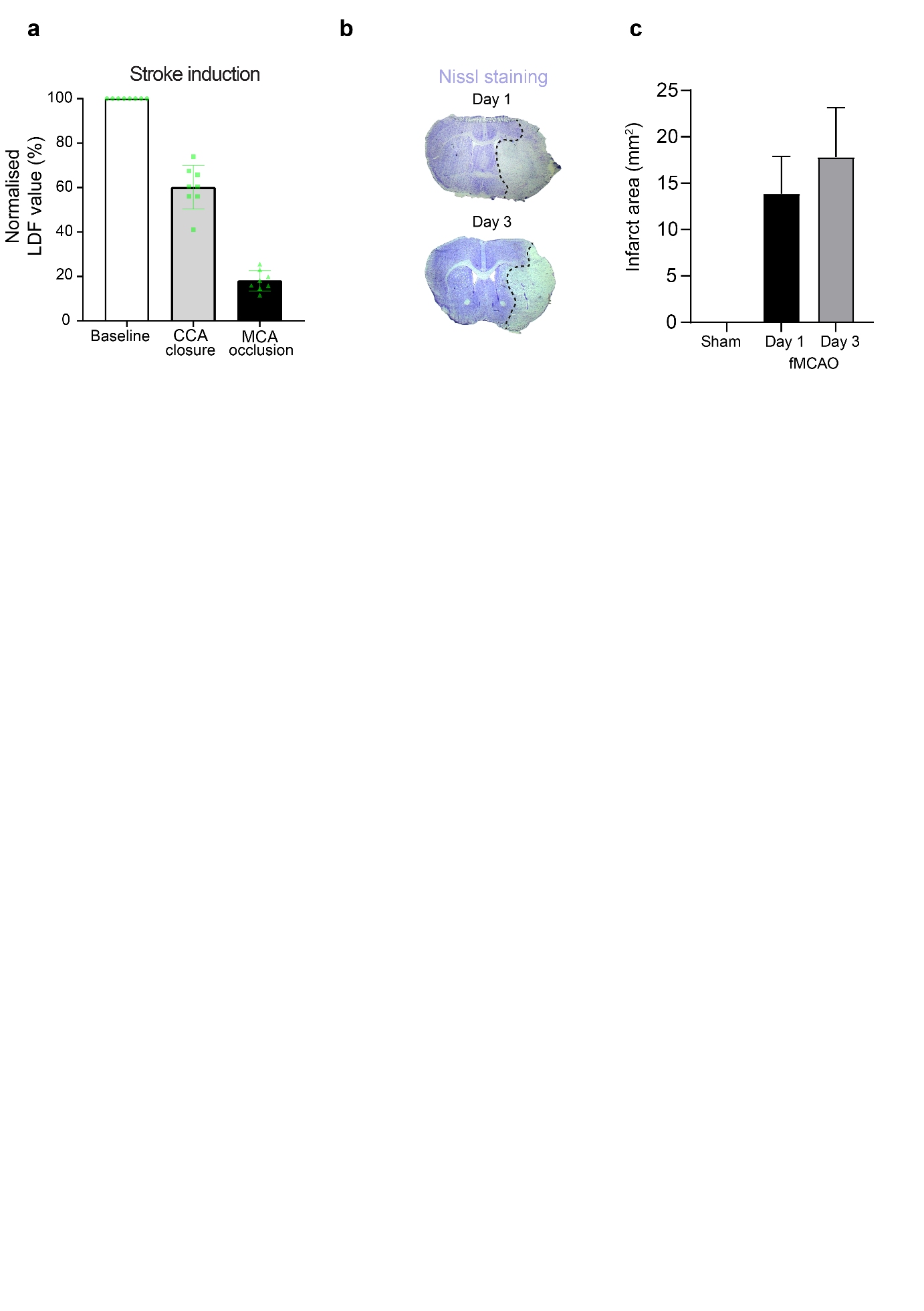

#### Supplementary Figure 7 Transient one-hour filament middle cerebral artery occlusion confirmation and loss of Nissl bodies indicating neuronal loss 24 hours and 3 days post stroke.

**a)** Stroke induction measured via laser speckle fluxmetry on the lateral temporal lobe of the skull placed to capture blood flow from the distal MCA territory. **b)** Nissl staining of the MCA territory of stroked animals on day 1 and day 3 post stroke (dotted lines indicate infarct core). **c)** Infarct area measured over the MCA territory on day 1 and day 3 post stroke. n= 5 adult male C57/Bl6/J mice 6-12 weeks old. Data is shown as mean +/- s.d.

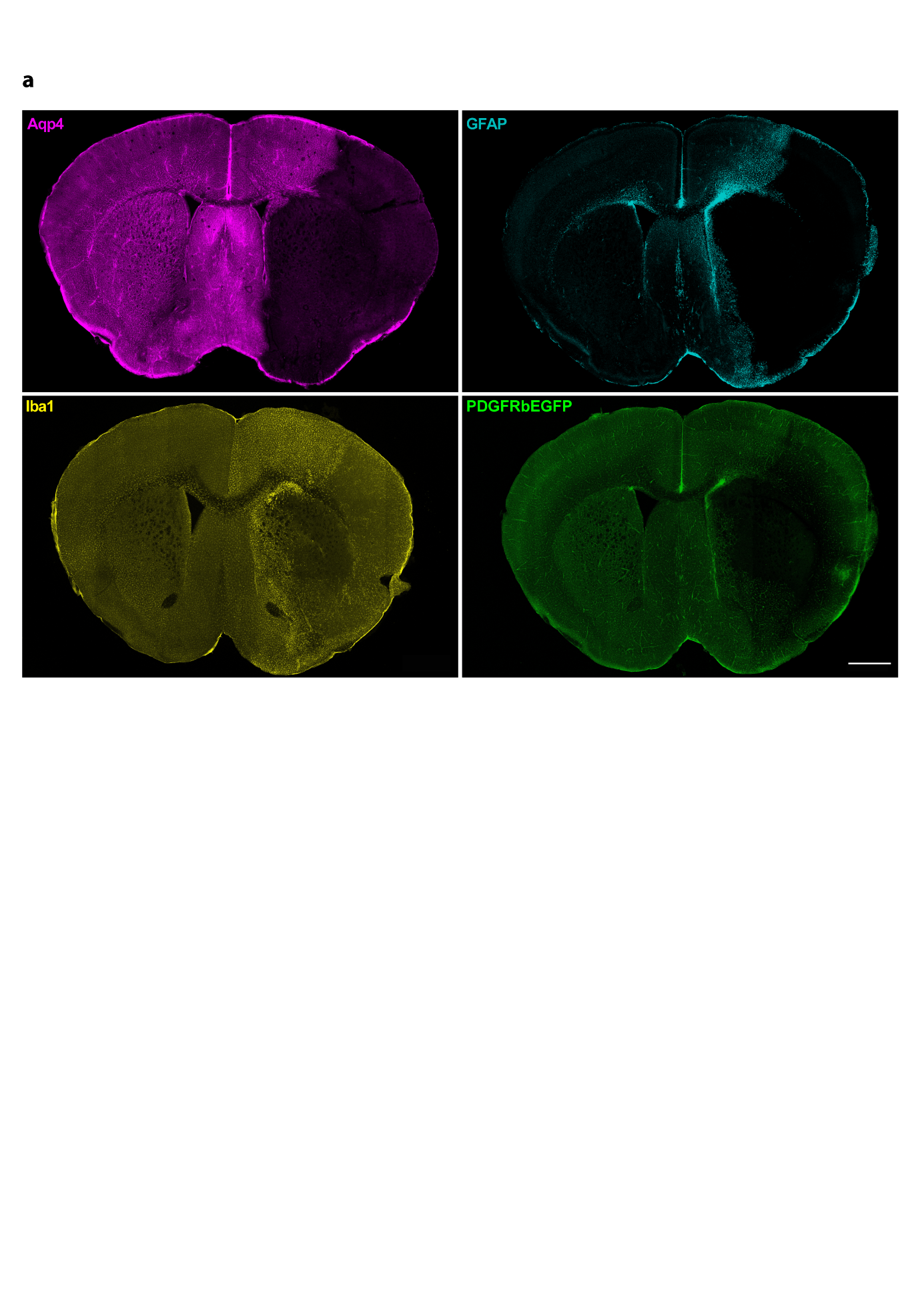

#### Supplementary Figure 8 Astrocyte, Microglial and mural cell distribution day 3 post-stroke.

**a)** Confocal overviews of immunostaining for aquaporin 4 (Aqp4), Glial fibrally acid protein (GFAP), iba1 (microglia) and PDGFRßEGFP in mice day 3 post stroke. Scale bars 1 mm.

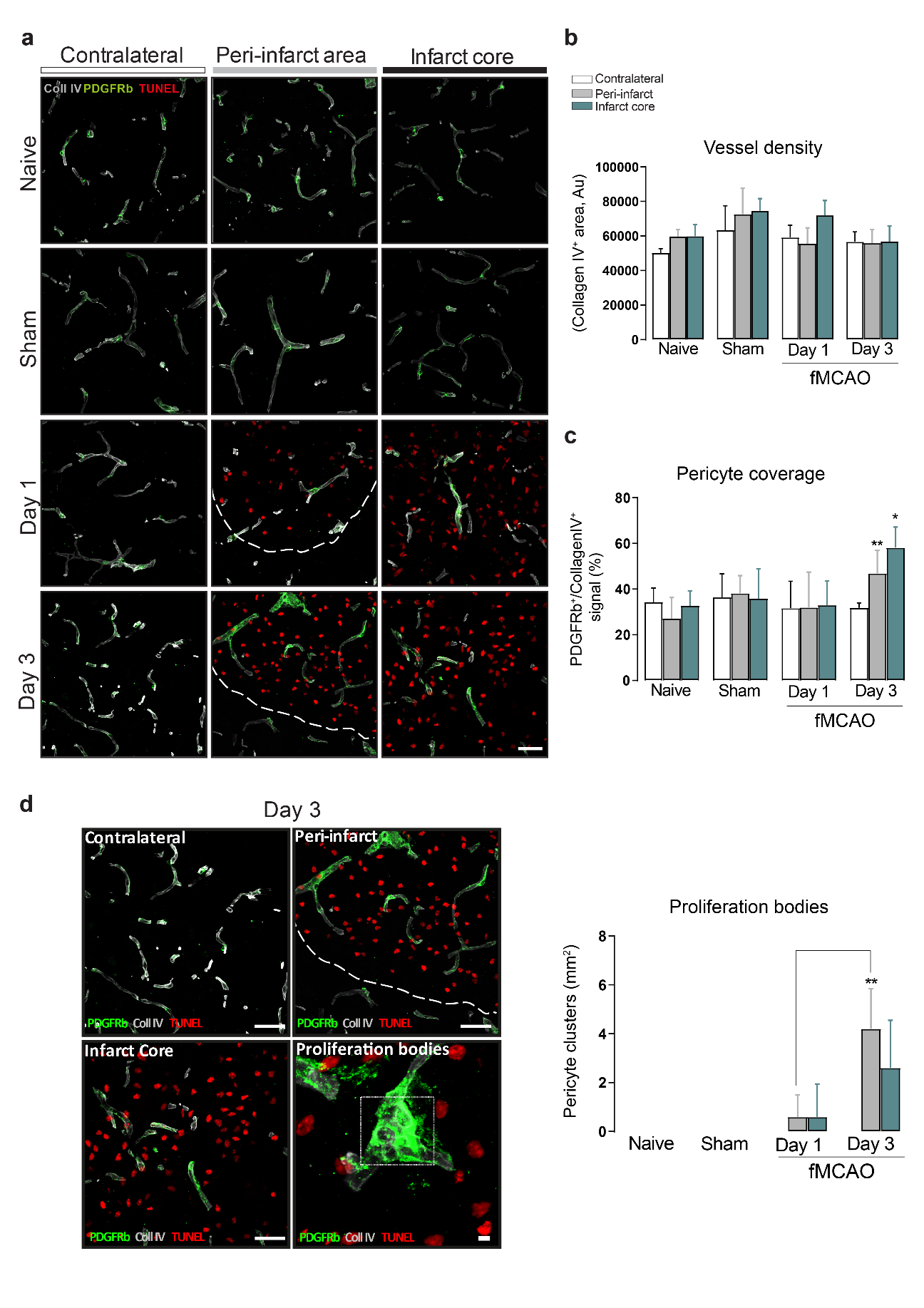

#### Supplementary Figure 9 Surviving pericytes expand vessel coverage and form clusters suggestive of proliferation on day 3 post stroke (24 hours – 3 Days post stroke).

**a)** Confocal micrographs of Collagen IV (gray), PDGFRß (green) TUNEL (red) immunofluorescence staining in C57BL/6J mice after no surgery, sham, day 1 or day 3 post stroke. **b)** Vessel density expressed as the amount of total pixels covered by Collagen IV+ pixels in each experimental group. **c)** Quantification of pericyte coverage expression as a ratio of PDGFRß+/Collagen IV+ pixel area. **d)** Confocal micrographs of the contralateral hemisphere and infarct core on day 3 post stroke respectively (left). Border zone of the infarct with the presence of PDGFRß+ and Collagen IV+ cell clusters within the infarct border zone (Right). Quantification of the appearance of proliferation bodies per mm2 induced by stroke. *P < 0.05, **P < 0.01, ***P <  0.005. n=5 mice per group, Statistics, unpaired two-tailed Student’s t-test, Scale bars, **a,** 50 µm and 10 µm, **d,** 50 µm and 5 µm. Data is shown as mean +/- s.d.

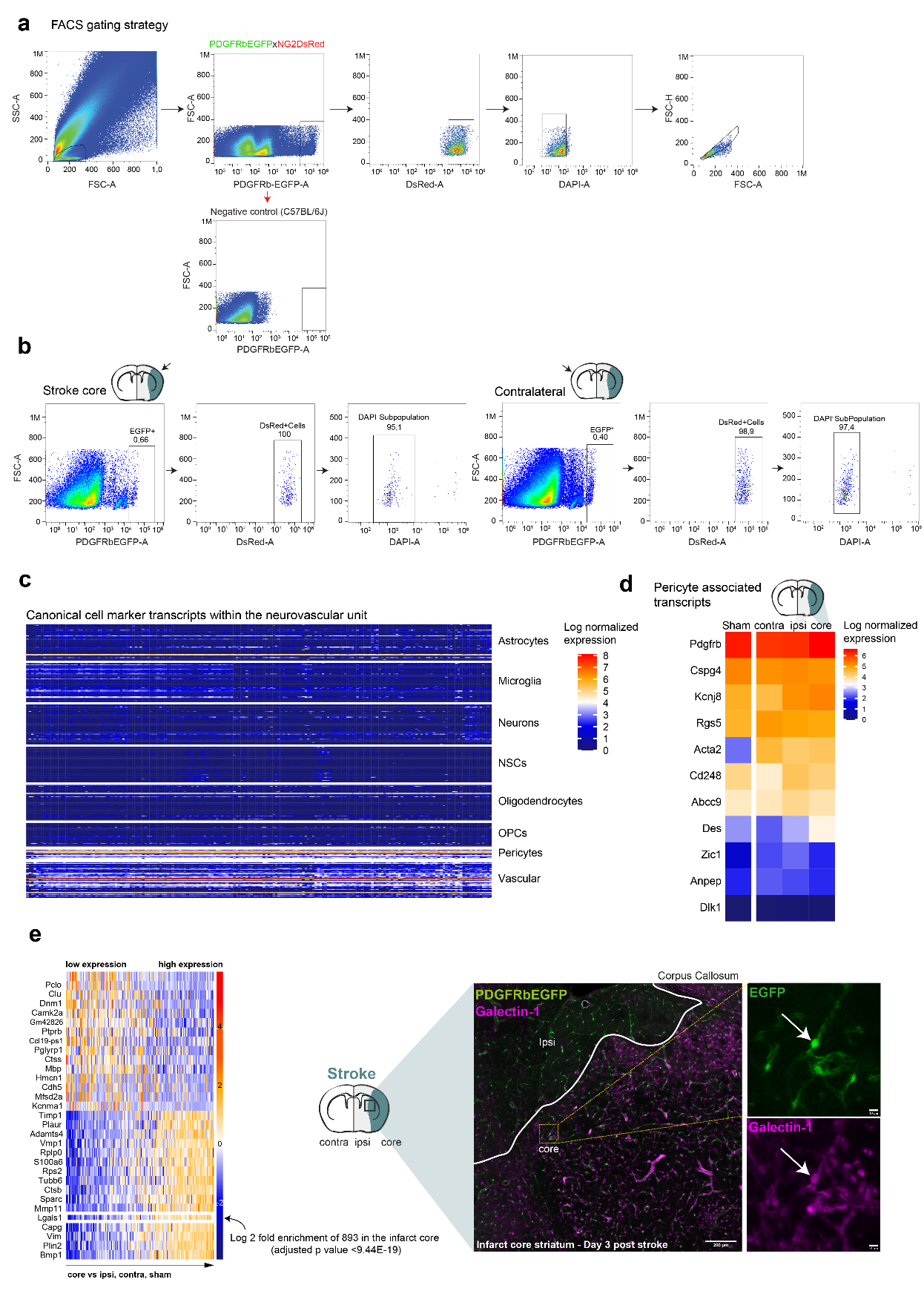

#### Supplementary Figure 10 RNAseq of pericytes within isolated brain regions identifies novel differential gene expression in ischemic pericytes on day 3 post stroke.

**a, b)** FACS gating strategy for isolation of EGFP/DsRED^+^ cell populations within core, ipsilateral and contralateral regions of day 3 post-stroked brains, including sham animals. **c)** Left, bulk RNA sequencing of canonical NVU cell population transcript markers. **d)** Enrichment of pericyte associated transcripts within the bulk sequencing data. **e)** Left, Differential gene expression of EGFP^+^/DsRed^+^ isolated mural cells shows enrichment or depletion of transcripts within the infarct core, compared to ipsilateral, contralateral and sham samples. Specifically, a large enrichment of lgals1 coding for the protein Galectin-1 was noted within the infarct core compared to other brain regions. Right, immunofluorescence microscopy of the Galectin-1 expression within the infarcted striatum of a PDGFRßEGFP mouse day 3 post stroke, validating the bulkRNA seq target. Scale bars, **e,** 200 µm and 10 µm

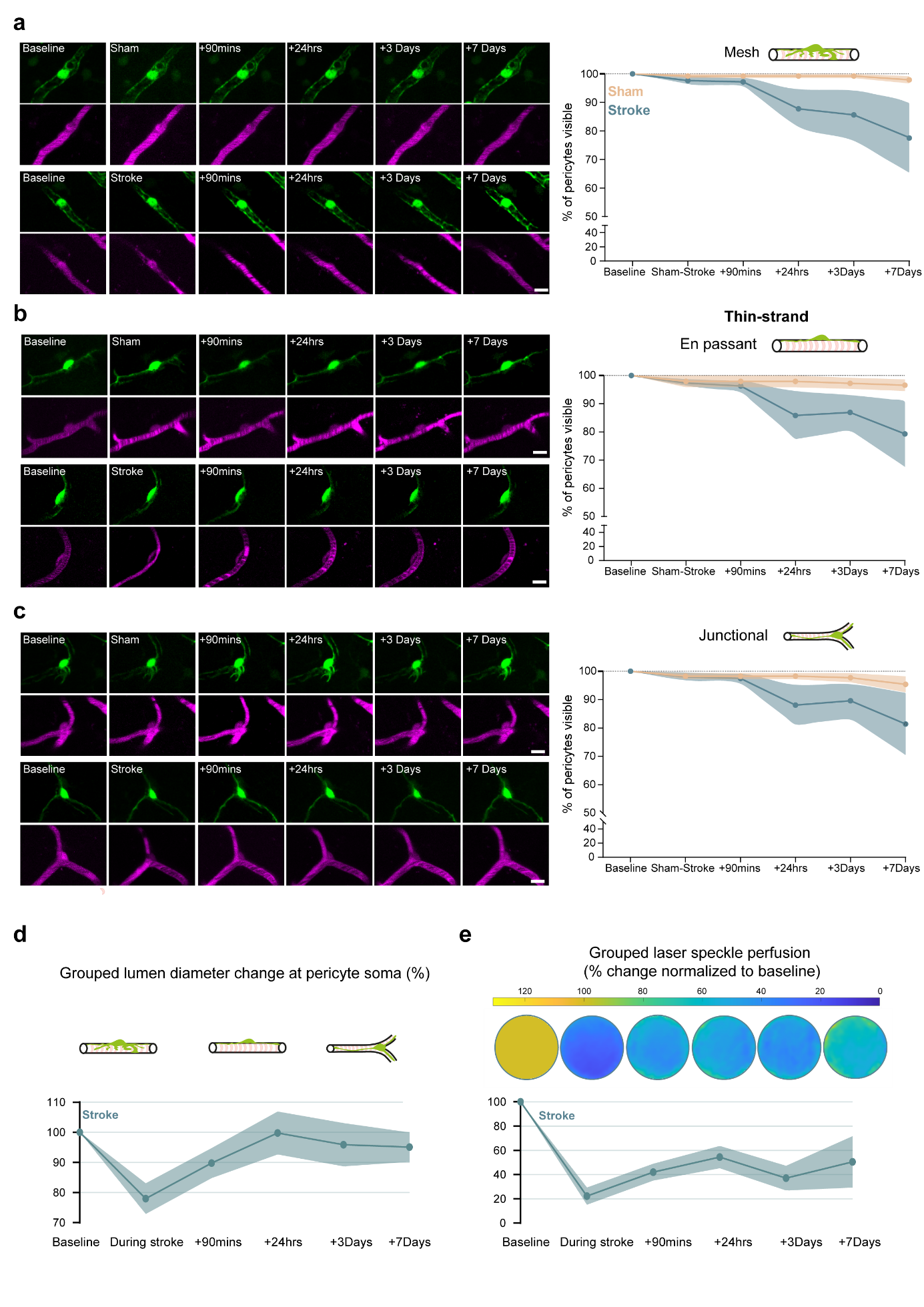

#### Supplementary Figure 11 Pericyte subtype loss, grouped capillary lumen diameter change and blood flow measurement within stroke cohort.

**a,b,c)** Left, individual pericyte subtypes per group (mesh, en passant and junctional thin-strand) over experimental timeline, green indicates pericytes and magenta indicates vascular flow. Right, pericyte subtype visibility over experimental timeline. **d)** Grouped lumen diameter change at pericyte soma within the stroked cohort. **e)** Upper, quantitative merging of all laser speckle data within stroked cohort per time-point pixel matched and normalized to baseline. Lower, normalized laser speckle perfusion normalized as a percentage to baseline. n=7 stroke mice and 4 sham mice ~1400 pericytes stroke-890 sham-510 across 6 timepoints. Scale bars, 5 µm. Data is shown as mean +/- s.e.m for **a,b,c** and s.d for **d** and **e**.

**Supplementary methods**

**Neuroscore and experimental stroke scale**

Following surgery mice were assessed for various neurological deficits, behavioral condition and core physiological parameters collectively named as the Neuroscore; which permits evaluation of each animal and determines termination end-points based on humane criteria. Experimental stroke scale (previously described here ^1^) was used as an evaluation to determine focal and general components relating to the mouse condition.

**FACS isolation of pericytes**

After three days of either stroke or sham surgery mice were sacrificed via cervical dislocation, brains were extracted and placed on ice cold 2% FBS in 1XPBS solution in a petri-dish. Brains were transferred to a brain mold and the cortex and striatum were dissected into three distinct regions: infarct core, ipsilateral hemisphere and contralateral hemisphere and placed in three separate petri dishes containing 3 ml of tissue collection solution (HBSS/BSA/glucose buffer) and 150 µl of DNAase stock solution at a concentration of 10 mg/ml. Brain tissue from each region was carefully dissociated mechanically using micro-forceps and placed in a 15 ml falcon containing tissue collection solution.

Falcon tubes containing the brain homogenate were centrifuged at 300 g at 4°C for 5 minutes. Supernatant was then discarded and 7.75 ml of pre-warmed collagenase/dispase solution (250 µl of collagenase/dispase 100 mg/ml stock solution in ultrapure dH_2_0 added to 7.5 ml of 2% FBS in 1XPBS) was added and the homogenate was resuspended and incubated on a rotisserie device in at incubator at 37°C for 30 minutes.

Falcon tubes were then centrifuged at 300 g at 4°C and the remaining cell pellet was resuspended in 1 ml of trituration solution (2% FBS in 1XPBS). Cell pellet was triturated carefully with p1000 pipettes, then p200 pipettes 100x respectively. The triturated cell solution was then centrifuged at 300 g for 5 minutes at 4°C and the pellet was resuspended in 1 ml of HBSS/BSA/Glucose buffer containing 40 µM of actinomycin and filtered through a 40 µm cell strainer and split into three FACS tubes. Tubes were then centrifuged at 300 g for 5 mins at 4°C and the supernatant was discarded, the resultant pellet was resuspended in 1 ml of HBSS/BSA/Glucose buffer containing 40 µM actinomycin, and immediately placed on ice for further processing. Each region: infarct core, ipsilateral hemisphere and contralateral hemisphere cell solutions were split into three tubes respectively each containing 1 ml of cell suspension in HBSS/BSA/Glucose buffer.

The first tube in each sample served to set up gating strategies for EGFP and dsRed while the second tube was incubated with DAPI at a concentration of 1:500 on ice for 20 minutes and was used to define the gating strategy for DAPI^-^ negative EGFP^+^ and DsRed^+^ cell populations. Finally, the third tube of each brain region: infarct core, ipsilateral hemisphere and contralateral hemisphere were used to sort cells for bulk sorting (50 cells in each well of a 96 well plate on a high yield setting) using the Sony SH800 cell sorter. This experiment was repeated 3X (with sham and stroke animals operated on in pairs).

**Library preparation for Smart-seq2**

The 96-well plates containing the sorted pools were first thawed and then incubated for 3 min at 72°C and thereafter immediately placed on ice. To perform reverse transcription (RT), we added to each well a mix of 0.59 μL H2O, 0.5 μL SMARTScribe™ Reverse Transcriptase (Clontech), 2 μL 5x First Strand buffer, 0.25 μL Recombinant RNase Inhibitor (Clontech), 2 μL Betaine (5 M Sigma), 0.5 μL DTT (100 mM), 0.06 μL MgCl2 (1 M Sigma), 0.1 μL Template-switching oligos (TSO) (100 μM AAGCAGTGGTATCAACGCAGAGTACrGrG+G). Next, RT reaction mixes were incubated at 42°C for 90 min followed by 70°C for 5 min and 10 cycles of 50°C 2 min, 42°C 2 min; finally ending with 70°C for 5 min for enzyme inactivation. Pre-amplification of cDNA was performed by adding 12.5 μL KAPA HiFi Hotstart 2x (KAPA Biosystems), 2.138 μL H2O, 0.25 μL ISPCR primers (10 μM, 5′ AAGCAGTGGTATCAACGCAGAGT-3), 0.1125 μL Lambda Exonuclease under the following conditions: 37°C for 30 min, 95°C for 3 min, 19 cycles of (98°C for 20 sec, 67°C for 15 sec, 72°C for 4 min), and a final extension at 72°C for 5 min. Libraries were then cleaned using AMPure bead (Beckman-Coulter) cleanup at a 0.7:1 ratio of beads to PCR product. Library was assessed by Bio-analyzer (Agilent 2100), using the High Sensitivity DNA analysis kit, and also fluorometrically using Qubit’s DNA HS assay kits and a Qubit 4.0 Fluorometer (Invitrogen, LifeTechnologies) to measure the concentrations. Samples were normalized to 160 pg/µL. Sequencing libraries were constructed by using an in-house produced Tn5 transposase (Picelli et al., 2014). Libraries were barcoded with the Illumina Nextera XT (FC-131-1096, Illumina) and pooled, then underwent three rounds of AMPure bead (Beckman-Coulter) cleanup at a 0.8:1 ratio of beads to library. Libraries were sequenced 2x100 reads base pairs (bp) paired-end on Illumina HiSeq4000.

**Analyses of Smart-seq2 data (Seurat cell cycle and Gene ontology using Metascape)**

BCL files were demultiplexed with the bcl2fastq software from Illumina. After quality-control with FastQC, reads were aligned using rnaSTAR ^2^ to the GRCm38 (mm10) genome with ERCC synthetic RNA added. Read counts were collected using the parameter “quantMode GeneCounts” of rnaSTAR and using the unstranded values. From that point, Seurat R v.3.1.2 package was used ^3^. Low-quality samples were filtered out from the dataset based on a threshold for the number of genes detected (min 1000 unique genes/pool), percentage of mitochondrial genes (max 0.75%), percentage of ERCCs (2.5% max) and number of reads (between 200k to 2M). 189 pools passed the quality-control. Gene expressions were log normalized using the NormalizeData function of Seurat with a scale factor of 100,000. Dataset were scaled and percentage of ERCCs and plate batches were regressed using ScaleData function. The top 2000 most variable genes were considered for the PCA. PC1 explained 4.4% and PC2 explained 1.8% of the variance. The pericyte score per sample is the average expression of the pericytes-specific genes from ^4^. The normalized counts were used to compute this average. Cell-cycle was assessed using the CellCycleScoring function and the cc.genes list provided in Seurat. Gene ontology analyses were performed using Metascape ^5^ by submitting the top head, (then top tail) 100 genes on the ordered PC loadings to interpret the principal components.

**Supplementary Table 1: Genes used for cell type identification of EGFP^+^DsRed^+^ isolated cells**

| genes used for cell type identification | | | | | | | |
| --- | --- | --- | --- | --- | --- | --- | --- |
| **Neurons** | **Astrocytes** | **Oligodendrocytes** | **Vascular** | **Microglia** | **OPC** | **NSC** | **Pericytes** |
| Trank1 | Gjb6 | Adamts4 | Eltd1 | Tmem119 | A930009A15Rik | Aurkb | Acta2 |
| Atp1a3 | Cldn10 | St18 | Abcb1a | Abi3 | C1ql1 | Cdca3 | Kcnj8 |
| Ptpn5 | Htra1 | Gjb1 | Ly6c1 | Olfml3 | Neu4 | Ckap2 | PDGFRß |
| Myt1l | Acsbg1 | Tmem125 | Flt1 | Mlxipl | Pdgfra | Ube2c | Cd248 |
| Scn2b | Ntsr2 | S1pr5 | Pglyrp1 | Apbb1ip | Nxph1 | Cdca2 | Anpep |
| Hpca | Slc7a10 | Gm98 | Cav1 | Selplg | Sema3d | Hmmr | Des |
| Gpr88 | Slc39a12 | Plxnb3 | Nostrin | Samsn1 | Shc4 | Tk1 | Dlk1 |
| Syt1 | Fgfr3 | 1700047M11Rik | Car4 | Il1a | Cspg4 | Pbk | Zic1 |
| Syt4 | AI464131 | Gal3st1 | Igfbp7 | Siglech | Chst5 | Neil3 | Abcc9 |
| Snap25 | Aqp4 | Cldn14 | Ly6a | Ccr5 | 2310050C09Rik | 1190002F15Rik | Rgs5 |
| Rab3c | F3 | Nipal4 | Ptprb | Trem2 | 8030423F21Rik | Casc5 | Cspg4 |
| Snca | Gpr37l1 | Gjc2 | Hspb1 | Gngt2 | Has2as | Kif4 |  |
| Pde10a | Mlc1 | Cyp2j12 | Itga1 | Cx3cr1 | Ltb4r1 | Bub1b |  |
| Rbfox1 | Bcan | Ppp1r14a | Cdh5 | Havcr2 | Mc5r | Ckap2l |  |
| Camk4 | Cyp4f15 | Sox10 | Icam2 | F11r | Olfr181 | Bub1 |  |
| Erc2 | Hepacam | Galnt6 | Akap12 | Itgam | Olfr869 | Ccnb1 |  |
| Celf5 | Fam107a | C030030A07Rik | E030010A14Rik | P2ry13 | Pnlip | Cdca5 |  |
| Snhg11 | Dio2 | Fa2h | Cd93 | Csf3r | Rdh16 | Kif2c |  |
| Slc12a5 | Rorb | Aspa | Clic5 | Susd3 | Gpr17 | Top2a |  |
| Camkv | Atp13a4 | Cdc42ep2 | Ly6c2 | Gpr34 | Matn4 | Ttk |  |
| Bcl11b | Ppap2b | Gjc3 | Higd1b | Parvg | Ppapdc1a | Depdc1a |  |
| Camk2b | Aldoc | Hhip | Tek | Mpeg1 | 2010317E24Rik | Ndc80 |  |
| Rasgef1a | Gja1 | Tmem88b | Wfdc1 | Bcl2a1b | Slc6a13 | Aspm |  |
| Gad2 | Sox9 | Opalin | Tbx3 | Mafb | Sox10 | Ect2 |  |
| Eno2 | S1pr1 | Padi2 | Gja4 | Tmem173 | Col12a1 | Ska1 |  |
| Gap43 | Slc6a11 | Mog | Esam | Ly86 | Gm7102 | Cdc25c |  |
| Grin2b | Slc1a3 | Cdk18 | Hmcn1 | Bcl2a1d | Emid1 | Troap |  |
| Scn2a1 | Ndrg2 | Insc | LOC545261 | Lair1 | Susd5 | Foxm1 |  |
| Gad1 | Nkain4 | Slc45a3 | Ctla2a | Ikzf1 | Col1a1 | E2f8 |  |
| Caly | Timp4 | Itgb4 | Tm4sf1 | Crybb1 | Mmp15 | Kif18b |  |
| AI593442 | Tril | D16Ertd472e | AU021092 | Slc2a5 | Tmem108 | Spc25 |  |
| Grm5 | Atp1a2 | Unc5b | Slco1a4 | Laptm5 | 4930447A16Rik | Birc5 |  |
| Sh2d5 | Pdpn | Lpar1 | Gpr116 | Cd53 | 4933422A05Rik | Cdc20 |  |
| Cpne5 | Ednrb | Car14 | S100a11 | P2ry12 | Art5 | Mis18bp1 |  |
| Rbfox3 | Fam176a | Bcas1 | Slc38a5 | Nlrp3 | Cyp2j8 | Cenpi |  |
| Ndrg4 | Gpc5 | Pllp | Ehd2 | Tgfbr1 | Lrrn4 | Cdk1 |  |
| Dlgap2 | Hapln1 | Ninj2 | Lama4 | Cyth4 | Npy | Psrc1 |  |
| Pde1b | 2900052N01Rik | Erbb3 | Arhgap29 | Hk2 | Hsd3b4 | Hist1h2ab |  |
| Cx3cl1 | Dbx2 | Ermn | Cldn5 | 4632428N05Rik | Muc5b | Prr11 |  |
| Cacna2d3 | Cyp2j9 | Prox1 | Osmr | Ctss | Tnfsf14 | Gm6531 |  |
| Nrsn2 | Bmpr1b | Pdlim2 | Aoc3 | C1qa | Olig2 | Trip13 |  |
| Celf4 | Ddah1 | Zfp536 | Tie1 | Cd37 | Olig1 | Rrm2 |  |
| Arpp21 | Slc1a2 | Olig1 | Cd97 | Aif1 | Xylt1 | Fam64a |  |
| Rgs9 | Aldh1l1 | Prr5l | Eng | Pld4 | Pcdh15 | E2f7 |  |
| Fbxl16 | Adrbk2 | Trim36 | Mustn1 | Ccl4 | Cys1 | Cep55 |  |
| Camk2a | Id2 | Nkx6-2 | Slc38a11 | C1qc | Lix1l | Spag5 |  |
| Rasgrp1 | Fzd2 | Plekhh1 | Abcg2 | AF251705 | Scrg1 | Hist1h2ao |  |
| Ptprn | Lhx2 | Hapln2 | Mylk | Ecscr | Pllp | Dlx1as |  |
| Chn1 | Scara3 | Nkain1 | Fn1 | Fyb | Enpp6 | Gtse1 |  |
| Pcp4 | Atp1b2 | Tnfaip6 | Cxcl12 | Il10ra | Inhbb | Esco2 |  |

**Supplementary Table 2: Cell cycle associated genes used in the Seurat cell cycle analysis**

| **G2/M genes** | **S genes** |
| --- | --- |
| Hmgb2,Cdk1,Nusap1,Ube2c,Birc5,Tpx2,Top2a,Ndc80,Cks2,Nuf2,Cks1b,Mki67,Tmpo,Cenpf,Tacc3,Fam64a,Smc4,Ccnb2,Ckap2l,Ckap2,Aurkb,Bub1,Kif11,Anp32e,Tubb4b,Gtse1,Kif20b,Hjurp,Cdca3,Hn1,Cdc20,Ttk,Cdc25c,Kif2c,Rangap1,Ncapd2,Dlgap5,Cdca2,Cdca8,Ect2,Kif23,Hmmr,Aurka,Psrc1,Anln,Lbr,Ckap5,Cenpe,Ctcf,Nek2,G2e3,Gas2l3,Cbx5,Cenpa | Mcm5,Pcna,Tyms,Fen1,Mcm2,Mcm4,Rrm1,Ung,Gins2,Mcm6,Cdca7,Dtl,Prim1,Uhrf1,Mlf1ip,Hells,Rfc2,Rpa2,Nasp,Rad51ap1,Gmnn,Wdr76,Slbp,Ccne2,Ubr7,Pold3,Msh2,Atad2,Rad51,Rrm2,Cdc45,Cdc6,Exo1,Tipin,Dscc1,Blm,Casp8ap2,Usp1,Clspn,Pola1,Chaf1b,Brip1,E2f8 |
